# Tracing the taxonomic distribution of plant cell wall degrading enzymes across the tree of life using feature architecture aware orthology assignments

**DOI:** 10.1101/2024.10.16.618745

**Authors:** Vinh Tran, Felix Langschied, Hannah Muelbaier, Julian Dosch, Freya Arthen, Miklos Balint, Ingo Ebersberger

**Affiliations:** Applied Bioinformatics Group, Inst. of Cell Biology and Neuroscience, Goethe University Frankfurt, 60438 Frankfurt am Main, Germany; Senckenberg Biodiversity and Climate Research Centre (BIK-F), Frankfurt, Germany; LOEWE Centre for Translational Biodiversity Genomics (TBG), Frankfurt, Germany; Institute of Insect Biotechnology, Justus-Liebig University, Giessen, Germany

**Keywords:** – Feature architecture, orthology, equivalog, ortholog search, phylogenetic profile, cell wall degradation, cellulase, pectinase

## Abstract

The decomposition of plant material is a key driver of the global carbon cycle, traditionally attributed to fungi and bacteria. However, some invertebrates also possess orthologs to bacterial or fungal cellulolytic enzymes, likely acquired via horizontal gene transfer. This reticulated mode of evolution necessitates ortholog searches in large taxon sets to comprehensively map the repertoire of plant cell wall degrading enzymes (PCDs) across the tree of life, a task surpassing capacities of current software. Here, we use fDOG, a novel profile-based ortholog search tool to trace 235 potential PCDs across more than 18,000 taxa. fDOG allows to start the ortholog search from a single protein sequence as a seed, it performs on par with state-of-the-art software that require the comparison of entire proteomes, and it is unique in routinely scoring protein feature architecture differences between the seed protein and its orthologs. Visualizing the presence-absence patterns of PCD orthologs using a UMAP highlights taxa where recent changes in the enzyme repertoire indicate a change in lifestyle. Three invertebrates have a particularly rich set of PCD orthologs encoded in their genome. Only few of the orthologs show differing protein feature architectures relative to the seed that suggest functional modifications. Thus, the corresponding species represent lineages within the invertebrates that may contribute to the global carbon cycle. This study shows how fDOG can be used to create a multi-scale view on the taxonomic distribution of a metabolic capacity that ranges from tree of life-wide surveys to individual feature architecture changes within a species.

## Introduction

Cellulose is the most common organic molecule on earth (Limayem and Ricke 2012). Per year, more than 100 billion tons of cellulose are produced in nature representing an enormous carbon sink. Most of this cellulose is deposited together with hemi-cellulose and lignin in primary plant cell (Field et al. 1998). Traditionally, it was thought that only certain bacteria and fungi produce the enzymes necessary to break down plant cell walls, e.g., (Post et al. 1990; Doi and Kosugi 2004; Zhao et al. 2014) resulting in the release of the bound carbon. Over the past years, evidence has accumulated that some invertebrate animals, such as mollusks and individual arthropods also encode intrinsic cellulolytic enzymes in their genomes (King et al. 2010; Busch et al. 2019; Muelbaier et al. 2024). Comprehensive surveys outside bacteria (Talamantes et al. 2016) are still missing, and consequently, the taxonomic distribution of plant cell wall degrading enzymes (PCDs) across the tree of life, and the repertoires of PCDs in individual species, is still largely unknown.

Currently, Biodiversity genomics initiatives are generating an overwhelming mass of genomic information from “non-model” organisms even from the remotest corners of the tree of life (Teeling *et al*. 2018; Rhie *et al*. 2021; Lewin *et al*. 2022; The Darwin Tree of Life Consortium 2022). This data provides a rich resource for investigating the taxonomic distribution and evolution of PCDs at an unprecedented scale and resolution. Tapping the potential of this resource necessitates tracing of proteins across large taxon collections (Pellegrini 2012a; Birikmen et al. 2021; Moi and Dessimoz 2023). The resulting presence/absence patterns, i.e., the phylogenetic profiles (PP) of the proteins, inform about when in evolutionary history the corresponding genes arose (Domazet-Lošo et al. 2007), in which species the associated function is likely represented (Pellegrini 2012b; Cantalapiedra et al. 2021), and if lineage specific duplications resulted in the extension of a gene family (Domazet-Lošo et al. 2007; Stolzer et al. 2012; Darby et al. 2017).

PPs come in two flavors. Those that are based on unidirectional searches (PP_US_) are populated with significantly similar proteins to a seed protein in other species. These are typically identified using software such as BLAST (Altschul et al., 1990), Diamond (Buchfink et al. 2021), or MMSEQ2 (Steinegger and Söding 2017). PP_OG_, on the other hand, capture the presence/absence pattern of orthologs to a seed protein across species, *i.e.,* of evolutionarily related proteins whose genetic lineages were separated due to a speciation event (Fitch 1970). The use of orthologs removes the ambiguity concerning the evolutionary relationships of genes subsumed in a PP allowing for a more precise reconstruction of the evolutionary past. Moreover, it tightens the link between the presence of a gene in an organism and of the expected function. This is because orthologs are considered the best guess when searching for functionally equivalent proteins across species (Dolinski and Botstein 2007; Fang et al. 2010; Gabaldón and Koonin 2013; Rogozin et al. 2014).

Unfortunately, the generation of PP_OG_ is computationally expensive (Sonnhammer et al. 2014). Even the fastest algorithms (Sahl et al. 2014; Kaduk and Sonnhammer 2017; Miller et al. 2019; Cosentino et al. 2024) cannot cope with the data masses released by the various biodiversity genomics projects (Langschied et al. 2024). Therefore, several databases provide access to pre-computed groups of orthologs across hundreds, or in individual cases few thousands of species (Li et al. 2003; Sonnhammer and Östlund 2015; Altenhoff et al. 2018; Mi et al. 2019; Howe et al. 2021; Zdobnov et al. 2021). While this data is an excellent starting point for generating phylogenetic profiles, an extension with further taxa is hard given the substantial run time complexity of the ortholog search. This substantially impairs the use of PP_OG_ in the biodiversity genomics era.

Targeted ortholog search tools ameliorate the computational burden of establishing PP_OGs_, (Ebersberger et al. 2009; Petersen et al. 2017; Aramaki et al. 2020; Feldbauer et al. 2020; Cantalapiedra et al. 2021; Rossier et al. 2021). As a common theme, they extend pre-compiled orthologous groups (core groups) with sequences from further taxa, and thus scale linearly with the number of taxa and genes investigated. Although powerful, the currently available algorithms share two main limitations. They require the existence of pre-compiled orthologous groups, and it is not possible to seed the ortholog search with a single sequence. Thus, they don’t offer a user experience that is comparable to that of popular tools like BLAST (Altschul et al. 1990). Consequently, the addition of add taxa to a PP_OG_ is straightforward, but adding custom gene sets is not. The second limitation is shared by all currently available ortholog search tools: The comparison of orthologs is limited to assessing their amino acid sequence similarities. Although individual tools consider protein features (e.g., Aramaki *et al*. 2020; Cantalapiedra *et al*. 2021), they do not score differences of protein feature architectures between orthologs. However, this information is invaluable for investigating functional divergence of orthologs (Nichio et al. 2017; Dosch et al. 2023).

Here, we present fDOG, the first software that integrates a profile-based, targeted ortholog search starting from an individual protein sequence with an assessment of the feature architecture similarities between the seed protein and its orthologs. We benchmark the performance of fDOG in establishing feature-architecture-aware PP_OG_ for large collections of proteins and taxa and compare it with state-of-the-art ortholog search tools. We then use fDOG to scan more than 18,000 species distributed across the tree of life for their PCD repertoires. Adopting uniform manifold approximation and projection (UMAP) as a novel way to visualize the resulting taxon-gene matrix, we provide an overview of the taxonomic distribution of plant cell wall degrading enzymes with unprecedented resolution. We show that lineage-specific changes in the enzyme repertoire coincide with a change in lifestyle and provide novel insights into the potential capacities of individual invertebrates to degrade plant cell walls.

## Results

### Workflow

fDOG performs a profile-based targeted ortholog search (Ebersberger et al. 2009). In brief, a profile Hidden Markov Model (pHMM) is iteratively trained with an increasingly large set of orthologous sequences (core orthologs). The final pHMM is then used to identify significantly similar candidate sequences in a target species. Each hit sequence serves as a query in a sequence-similarity based search against a proteome of a reference species that is represented in the core ortholog group. The candidate is propagated to an ortholog if it either identifies the core ortholog as a best hit, or if the reference protein is a lower ranking hit, but its evolutionary distance to the best hit is smaller than the distance between the ortholog candidate and the best hit. If an ortholog relationship is established, fDOG computes, by default, the feature architecture similarity (Dosch et al. 2023) between the seed protein and the ortholog. The main workflow is summarized in Figure 1 and the full workflow is provided as Fig. S1. In the following sections, we describe the individual steps of fDOG.

**Figure 1.**
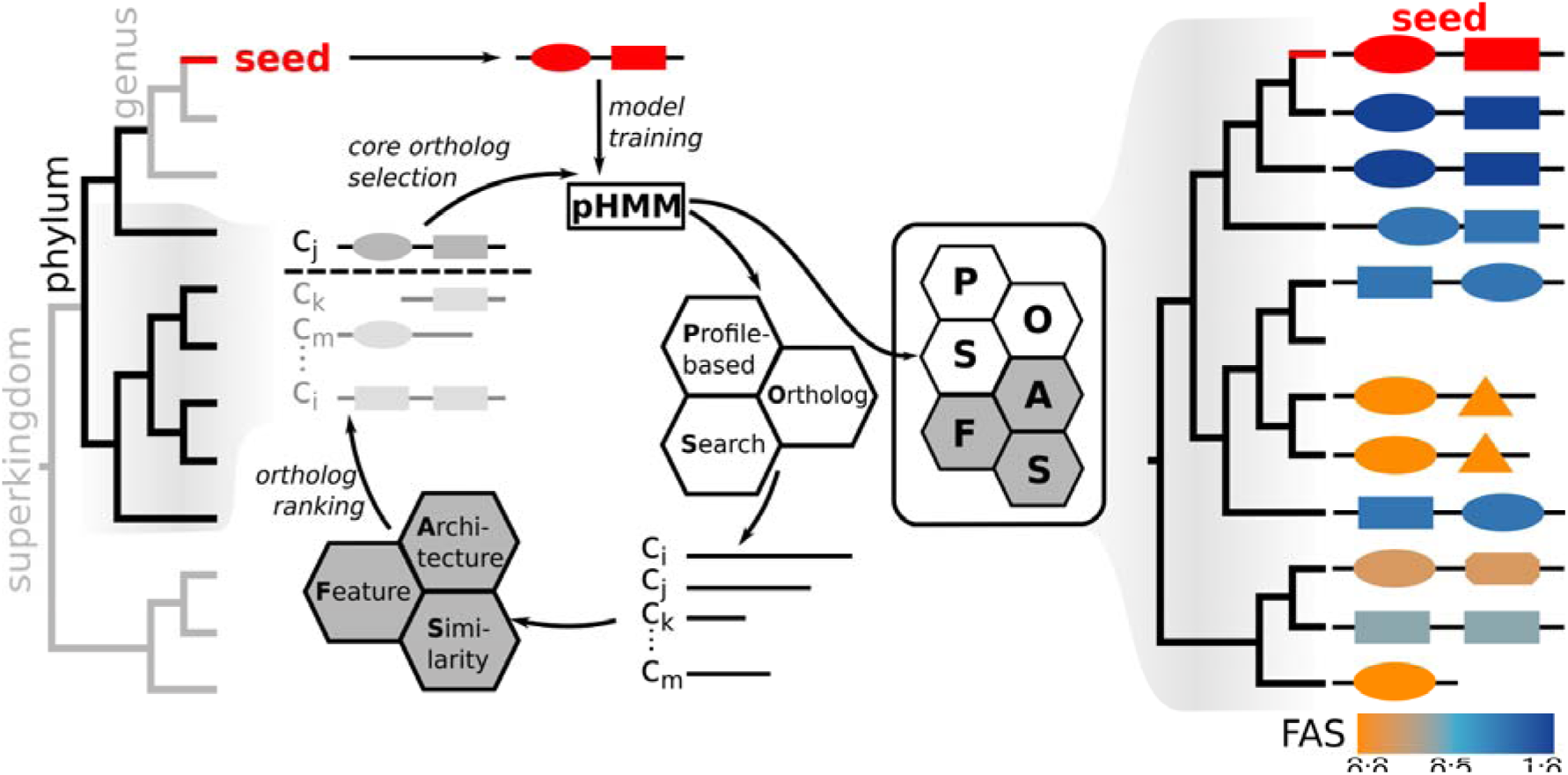
Feature-aware phylogenetic profiling with fDOG. pHMM Training phase: fDOG extracts the core orthologs used for pHMM training from a set of primer taxa that are ordered according to increasing taxonomic distance to the seed taxon from which the seed protein was taken from (left tree). A minimum and a maximum taxonomic distance of the primer taxa from the seed taxon can be specified to control the diversity of the training data (subtree in black). During pHMM training, fDOG iteratively compiles the core ortholog group and re-trains the pHMM. In each iteration, fDOG uses the current pHMM to identify orthologs across the primer taxa (*C_i_,…,C_m_*), it infers their feature architecture similarities (FAS) to the seed protein and orders them according to decreasing similarity of both amino acid sequence and feature architecture to the seed. The highest-ranking sequence is added to the core ortholog set. At the end of each iteration, the core orthologs are aligned and the pHMM is re-trained. Full ortholog search: The final pHMM is used for a profile-based ortholog search across all taxa from a proteome collection. The feature architecture of each detected ortholog is compared to that of the seed protein, and a similarity score (FAS score) is computed. This obtains the feature architecture aware phylogenetic profile of the seed protein (right tree). The color represents the FAS score ranging from 0 (orange; no shared features) to1 (blue; identical feature architectures). See main text for further information. The full workflow is shown in Fig. S1.

### Dynamic core ortholog compilation

fDOG starts the profile-based ortholog search across *n* taxa with a single protein sequence, the *seed*. We define the set *C* of core orthologs that are used for the pHMM training, the set of primer species *P* from which the core orthologs have been identified, and the set of search species *S*. We initialize *C* with the seed sequence *c_0_*, *P* with the species *p_0_* from which *c_0_* was taken, and *S* with the remaining n-1 taxa. Additionally, *T_S_* represents the NCBI common tree spanning the taxa in *S*. *pHMM_1_* is then trained with the sequence in *C*. To control the taxonomic diversity within the core ortholog group, fDOG can prune all taxa from *T_S_* below a minimal (option *--minDist;* default ‘genus’) or above a maximal (option *--maxDist;* default ‘superkingdom’) taxonomic distance from *p_0_*. Moreover, fDOG provides the option to specify a sub-set of *S* from which the core orthologs should be derived (option *--coreTaxa*). In each of the following *i* iterations, fDOG then traverses *T_S_* in the order of increasing taxonomic distance to *p_0_* where equidistant taxa are visited in random order. In each taxon, a targeted ortholog search using pHMM_i_ is performed (Ebersberger et al. 2009). We then add the ortholog *c_i_* to *C* and the corresponding *p_i_* to *P* that maximizes

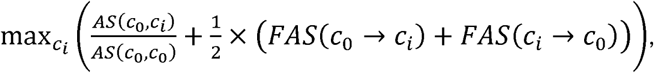

where *As*(*c*_0_,*c_i_*) is the pair-wise alignment score between ci and the seed protein, and *As*(*c*_0_,*c*_0_) is the maximally achievable alignment score. *FAS*(*c*_0_ → *c_i_*) and *FAS*(*c_i_* → *c*_0_) are the feature architecture similarities between the seed protein and the ortholog using either the seed or the ortholog as the reference. Ties are broken by selecting the sequence with the smallest taxonomic distance. The sequences in *C* are aligned with MAFFT-LINSI (Katoh and Standley 2013), and the *pHMM_i_* is re-trained. The iteration is concluded by pruning *p_i_* together with all species with a taxonomic distance below *minDist* to either *p_0_* or *p_i_* from *T_S_*. This has the effect that the taxonomic diversity of the core ortholog set either remains constant or increases with each iteration. We repeat the procedure unless (i) the specified number of core orthologs has been reached (option *--coreSize*), (ii) the leaf set of *T_S_* is empty, or (iii) no ortholog has been found in the last iteration.

To accelerate the core ortholog compilation, we devised two stopping rules: A protein is directly accepted as a core ortholog if (i) it reaches the maximally possible score of 2 (hard max score cut-off) or if (ii) it’s total score cannot be superseded by a user-specified margin (soft max score cut-off; default value 5%; set using the option *--distDeviation*) with a sequence from a taxonomically more distant clade in *T_P_*. Moreover, we skip the computationally costly FAS score calculation in cases where even the maximal FAS score of 1 would not suffice to make the ortholog a top-ranking candidate.

### Full ortholog search

In this phase of the *search*, fDOG uses the pHMM generated in the training phase—or from an existing pHMM library generated by a previous fDOG run—to identify orthologs across all taxa in *S*. Because there are no dependencies between the individual ortholog searches, this part of fDOG is parallelized. Each of *n* threads performs the ortholog search in a non-overlapping subset of *|S|/n* taxa. Without an impact on the performance of the ortholog identification, users can confine the search to individual taxonomic groups represented in *S.* It is also possible to later extend the search to include further taxa by re-using the pre-trained pHMM. A full list of options is provided in the software manual (see https://github.com/BIONF/fDOG/wiki/Options). Upon completion of this phase, fDOG outputs the identified orthologs to *c_0_* together with the taxa that harbor them in a tabular format. The corresponding amino acid sequences are stored in an accompanying FASTA file.

### Construction of feature architecture aware phylogenetic profiles

In the last *phase*, fDOG scores the feature architecture similarity (FAS) of each detected ortholog to the seed protein *c_0_*. Because FAS scores are not symmetric (Dosch et al. 2023), they are computed in both directions using in turns the seed protein (FAS_F) and the ortholog (FAS_B) as reference. The resulting scores are appended to the phylogenetic profiles, and the corresponding feature architectures are saved as an accessory output file in json format. In addition, a configuration file is produced for an easy upload of the phylogenetic profiles into PhyloProfile (Tran et al. 2018).

### Availability

fDOG is available from https://github.com/bionf/fdog under the GPLv3 license

#### Benchmark

fDOG generates phylogenetic profiles for custom sets of seed proteins across a user-defined collection of target species. By doing so, it conceptually differs from standard ortholog search tools that aim to identify orthology relationships for all proteins in a proteome. To still leverage the available framework for benchmarking ortholog search tools (Nevers *et al*., 2022), we computed the PP_OG_ for the 20,600 human proteins in the human reference proteome across the 77 further species in the Quest for Orthologs (QfO) reference proteome set.

### Core set compilation

We used 5 iterations to compile the core orthologs for each seed protein (see (Ebersberger et al. 2009)). After the training phase, 90% of the resulting core ortholog groups contained the full set of six sequences (see Fig. S2). For 494 human sequences, fDOG found no ortholog. These proteins are significantly shorter compared to proteins for which at least one ortholog was detected (Fig. S3). This is in line with a previous observation that it is harder to identify orthologs for short proteins (Jain et al. 2019). We next evaluated the orthology relationships for consistence with assignments made by the 17 orthology predictors available in the Quest for Ortholog Benchmark Service. Only 982 out of 97,234 predicted core ortholog pairs were exclusively predicted by fDOG (Fig. S4). These fDOG-only orthologs were found more frequently in species that are evolutionarily more distantly related to humans (Fig. S5). 38 core ortholog groups comprise only fDOG-only orthologs. To assess if these assignments are largely false positives, we computed maximum likelihood gene trees for the 20 groups with at least four proteins and compared the resulting tree to the species phylogeny. In all cases, the ML tree was either identical or not significantly different from the species tree (Table S1), providing no indication for a false positive orthology assignment.

### Profile-based targeted ortholog search

We next benchmarked the performance of fDOG in the final ortholog search, where the fully trained pHMM is used. We identified orthologs to the human proteins across all reference proteomes (Fig. S6), and investigated the performance of the ortholog search using the tests from the Orthology Benchmark Service (Nevers *et al*., 2022). We compared fDOG to InParanoid (Persson and Sonnhammer 2022) and OMA pairs (Zahn-Zabal et al. 2020), because similar to fDOG both tools compute orthologs between pairs of species. fDOG identified 579,197 pairwise orthologs to human proteins across the 77 species in the reference proteome set, which ranges between OMA (505,380) and InParanoid (639,992; Fig. S7). The results for three benchmark tests are summarized in Figure 2. This reveals that the use of the full set of ortholog assignments already results in a performance of fDOG that is comparable to that of state-of-the-art tools that require the analysis of entire proteomes. However, one of the main innovations of fDOG is the computation of the bi-directional feature architecture similarities between the orthologs (FAS score). This offers the flexibility of adjusting the stringency of the ortholog search *post hoc* via a dynamic filtering of the orthology assignments using the FAS score. To evaluate the impact of this filter on the benchmark, we removed fDOG ortholog assignments with a FAS score below 0.75 applying the filter either to the FAS_F, the FAS_B, or the mean of the two FAS scores (see Figs. S8 and S9 for other thresholds). This improved the quality metrics in all three tests surpassing the performance of the other two tools in individual cases.

**Figure 2.**
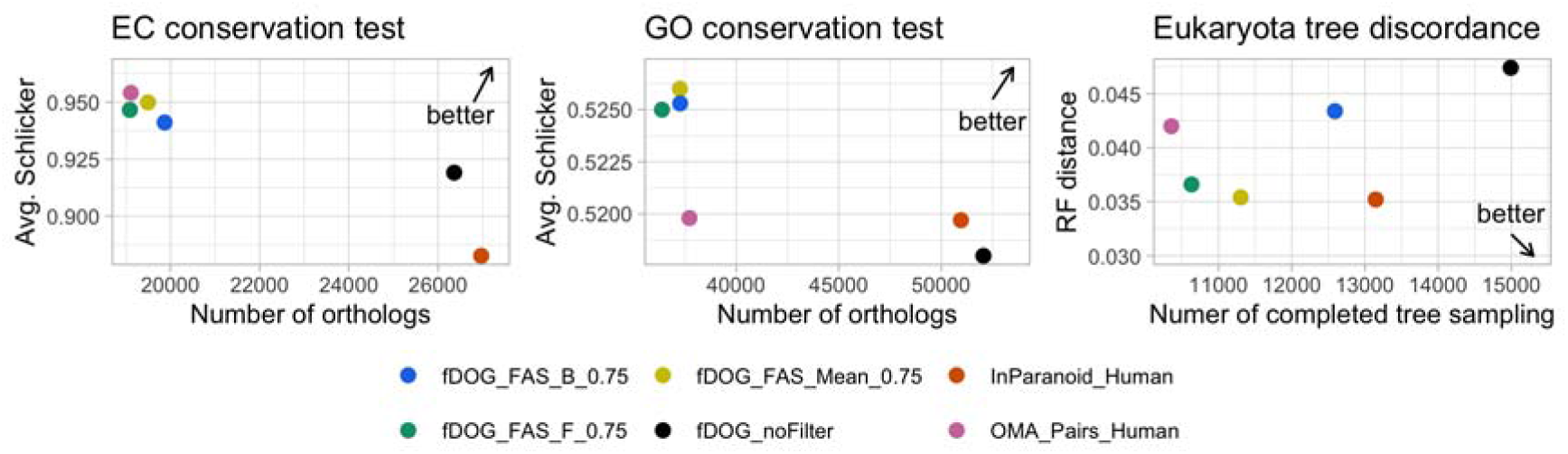
Benchmark of fDOG using three tests from the Quest for Orthologs benchmarking service (Nevers et al., 2022). fDOG ortholog assignments were either taken as is (fDOG_noFilter) or filtered to contain only ortholog pairs with a FAS score of at least 0.75 using either the seed protein as reference (FAS_F_0.75), the ortholog as reference (FAS_B_0.75), or the mean of the two FAS scores (FAS_mean_0.75). Each dot in the three plots indicates the test result for the respective method. The direction of the arrow in each plot indicates the direction where better results are to be found. The EC conservation test determines if orthologs of human enzymes are annotated with the same EC number. The GO conservation test computes the semantic similarity of the GO term annotation of the orthologs identified by the individual tools. The species tree discordance test compares the topology of the gene tree computed from the orthologs to that of the accepted species phylogeny. RF - Robinson-Foulds distance *(Robinson and Foulds 1981)*.

### Runtime

The computation of the feature architecture aware PP_OG_ for 20,600 human proteins across the 77 other taxa in the QfO 2020_04 reference dataset required 82 hours on a compute server with 2 processors AMD EPYC 7601 (32 cores; 2.2Ghz). The core ortholog compilation across all seed proteins was completed after 18 h, with an average of 183 seconds per protein. The subsequent targeted ortholog search across all search taxa required 48 h with an average of 8 seconds per protein (Figs. S10 and S11). The computation of the bi-directional FAS scores took 16h. As expected from the design of fDOG, the runtime of the ortholog search scales linearly with the number of search taxa (Fig. S12).

#### The taxonomic distribution of plant cell wall-degrading enzymes

fDOG is the first tool that can compile, without loss of performance, feature architecture aware PP_OG_ for individual genes of interest. We next used fDOG for tracing enzymes that are potentially involved in the degradation of plant cell walls across the tree of life. Using the CAZy database (cazy.org (Drula et al. 2022); last accessed 25.06.2024), we selected 61 enzyme sub-families whose activities indicate that they are associated with plant cell wall degradation. This included 42 cellulases, 9 hemi-cellulases and 10 pectinases (Table S2). We next identified a minimal set of seven seed species (3 fungi, 3 bacteria and 1 mollusk) such that each CAZy sub-family is represented by a protein in at least one of the seed species (Table 1). We then identified all proteins assigned to any of the 61 families in the seed species. We generated a non-redundant seed protein list by keeping for each sub-family only proteins from one of the seven species. Whenever possible, we gave preference to the plant pathogen *Rhizoctonia solani* (fungi). We refer to the resulting set of 235 enzymes as potentially plant cell wall degrading enzymes (pPCDs) (Tables 1 and S2). Each protein was then used as a seed for an fDOG search across 18,565 taxa (1,354 eukaryotes, 887 archaea and 16,324 bacteria).

**Table 1.**
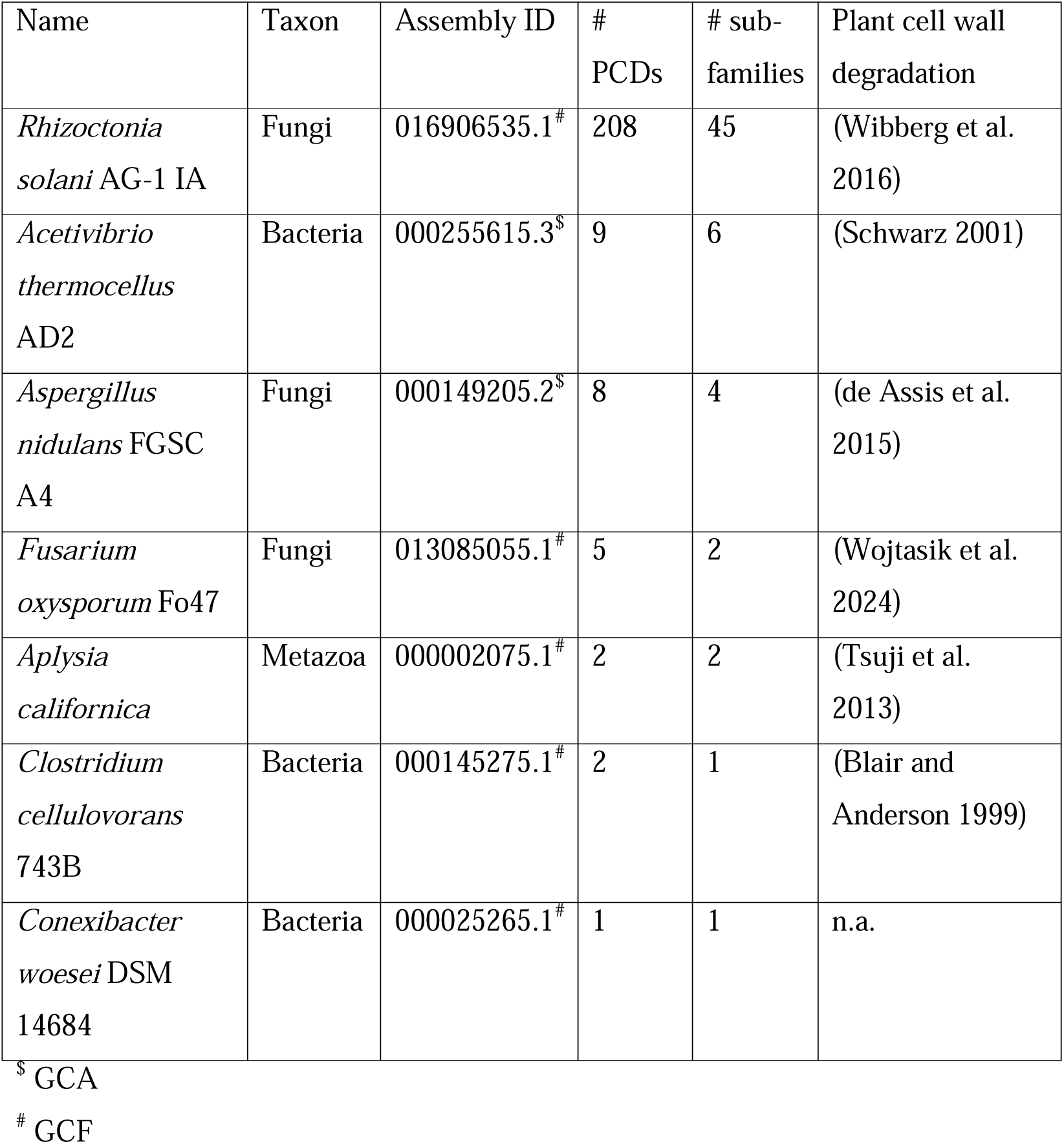
Seed species for the phylogenetic profiling of pPCDs

The resulting phylogenetic profiles are provided as Supplementary data 1. A preliminary analysis revealed that many of the fungal seed proteins are paralogs that arose by a gene duplication within the fungi. To make the analysis more concise, we selected for each group of fungal paralogs one representative (Table S2). The resulting reduced matrix comprises orthologs for 76 proteins and is shown in Fig. S13. With this number of genes and taxa investigated, a conventional taxon gene matrix provides little more than the overall distribution pattern of the genes. To provide a more accessible visualization, we extracted for each of the search taxa a binary vector encoding the presence or absence of the 76 representative proteins. We then used these vectors as input for a dimension reduction with UMAP (McInnes et al. 2020) (Fig. 3; for an interactive plot see Fig. S14). In the resulting plot, the topological placement of the individual species is determined by their repertoire of pPCDs. Interestingly, species from the same taxonomic group exhibit a tendency for spatial clustering. For example, Fig. 3A reveals that most fungi and plant species are placed in spatial proximity, respectively. This allows to identify similarities but also (unexpected) variations in the pPCD repertoire among closely related species. Within the bacteria, some taxa, e.g. the *Actinomycetota*, *Bacillota* and *Bacteroidota* are substantially more spread out than both plants and fungi. Some of these taxa are renowned for their cellulolytic capabilities (Doi and Kosugi 2004; Berlemont and Martiny 2013). In the UMAP, these are readily distinguished from other members of their clade by forming prominent clusters of species rich in pPCDs (see Fig. 3B for the *Actinomycetota*, and Fig. S14 for the *Bacillota* and *Bacteriodota*, respectively).

**Figure 3.**
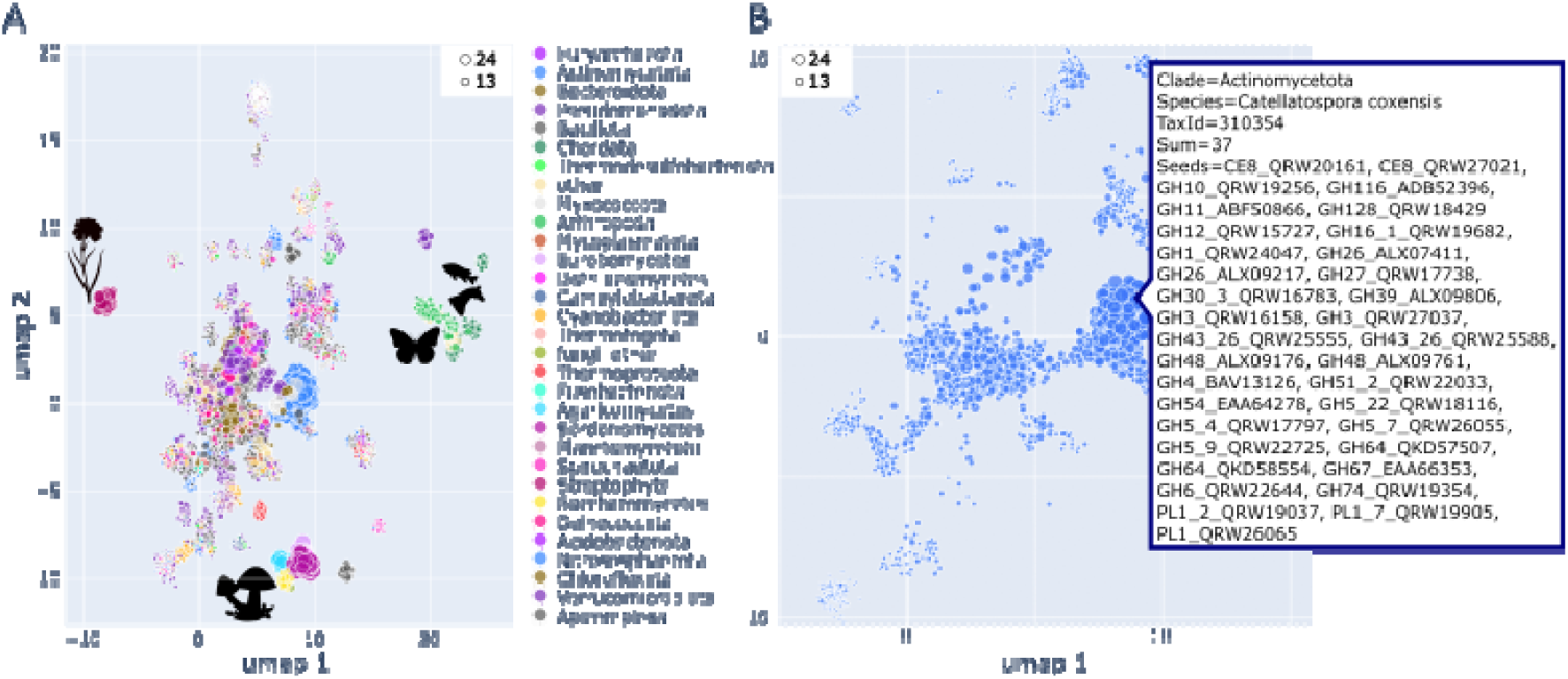
UMAP of 18,565 taxa from the three domains of life based on their pPCD repertoire. Each dot represents a species, and its placement in the plot is determined by the presence-absence pattern of orthologs to the 76 representative pPCDs. Dot sizes are proportional to the number of seed genes represented by at least one ortholog (see legend in the top corner of the plot). Selected taxonomic groups are indicated by color. A) All taxa. Pictograms indicate clusters that mainly contain plant, fungal and animal taxa. An interactive representation of this plot which provides the option to filter taxa, and to explore the pPCD sets for individual species is provided in Figure S14. B) Variation of pPCD repertoires across the Actinomycetota. The inlay provides detailed information for the soil bacterium *Catellatospora coxensis*, a bacterium with one of the largest and most diverse sets of pPCD orthologs in our analysis.

### Variation of pPCD repertoires indicate a change in lifestyle

To obtain a higher resolving picture, we next shifted our focus on eukaryotic species (Fig. 4A). This revealed that fungi are represented by two distinct clusters. One comprises species with smaller and less diverse pPCD repertoires. The second cluster represents species with richer pPCD repertoires. Individual fungal classes are placed exclusively in one of the two clusters, such as *Saccharomycetes* (Cluster 1), *Tremellomyetes* (Cluster 1) or *Agaricomycetes* (Cluster 1). This indicates that their pPCD repertoires were subject to little change during evolution. Other fungal classes, are, however, spread over both clusters. For example, most *Sordariomycetes* are placed in the fungal cluster 2, except for 11 species that are characterized by a reduced set of pPCDs indicating a loss of the corresponding genes (see Figs. 4A,B and Fig. S15). Interestingly, 9 of these 11 species parasitize either insects or nematodes (Table 2) implying that plant cell wall degrading capabilities are less important. A similar pattern can be observed in *Dothideomycetes*. Many members of this class are plant pathogens, endophytes or grow on decaying plant material. They are equipped with a rich set of pPCDs. In contrast, six species show a reduced set of pPCDs and are placed in cluster 1 (see Figs. 4A and B and Table 2). Among these are the “whiskey fungus” (*Baudoinia panamericana*), which uses volatile carbohydrates as energy source (Ewaze et al. 2007), and an extremophile black yeast (*Neohortea acidophila*) capable of using lignite as its sole carbon source (Hölker et al. 2004).

**Figure 4.**
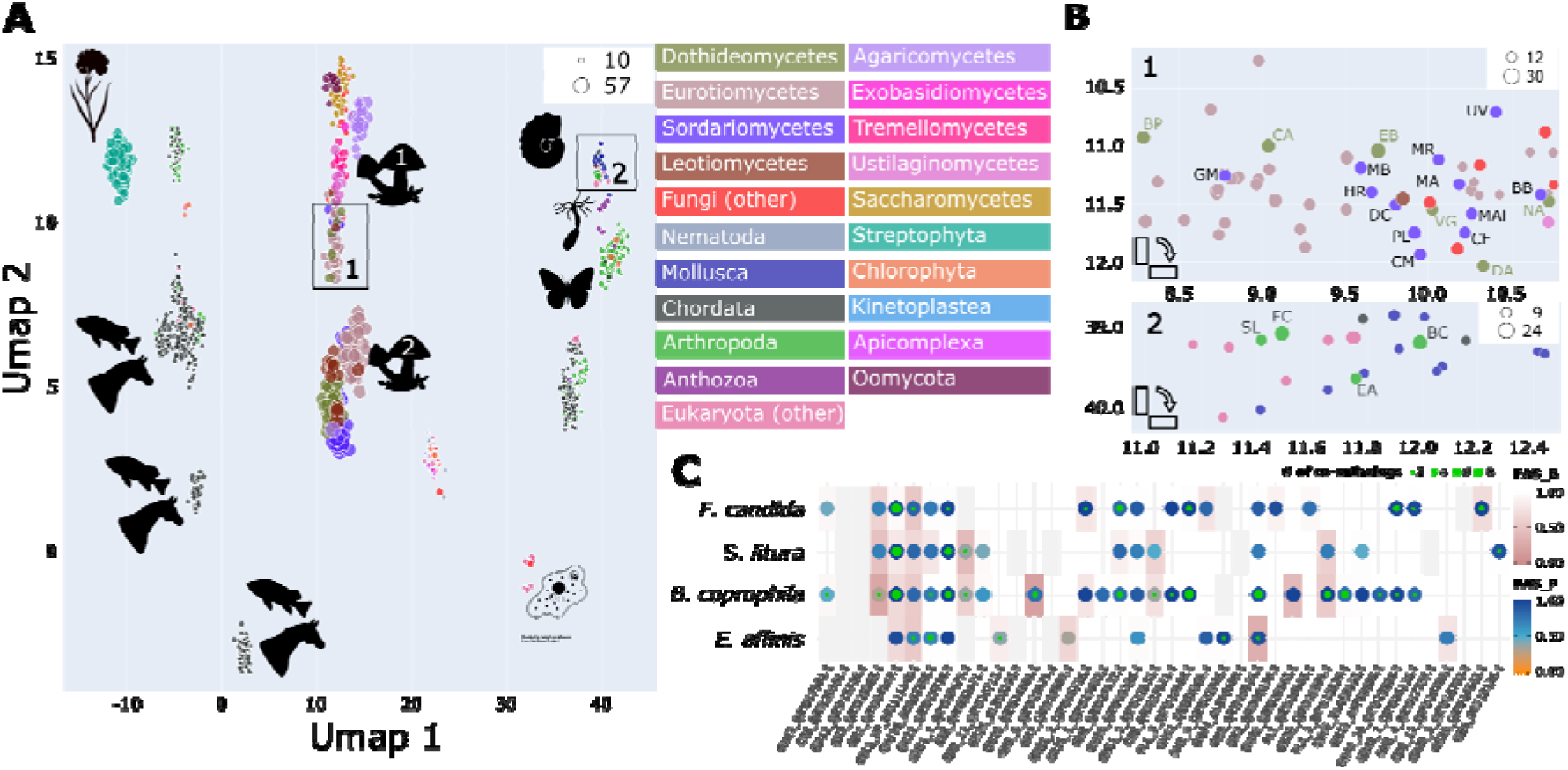
pPCD repertoires across the eukaryotes. A) UMAP summarizing information for all 1,354 investigated eukaryotic species. Each species is represented by a dot, and the spatial placement is determined by the pPCD repertoire. The dot size is proportional to the number of pPCDs with at least one ortholog in the representative species. Pictograms represent taxa that are preferentially found in the individual cluster. A zoom on the candidate regions 1 (fungal cluster 1) and 2 (invertebrate cluster) is shown in B). Plot layout and color coding follows A). Sordariomycetes (purple): GM – *Geosmithia morbida*; MB – *Metarhizium brunneum*; HR – *Hirsutella rhossiliensis*, DC – *Drechmeria coniospora*; CM – *Cordyceps militaris*; MR – *Metarhizium robertsii*; MA – *Metarhizium acridum*; MAl – *Metarhizium album*; CF – *Cordyceps fumosorosea*; UV – *Ustilaginoidea virens*; BB – *Beauveria bassiana; PL – Purpureocillium lilacinum.* Dothideomycetes (moss green): BP – *Baudoinia panamericana*; CA – *Coniosporium apollinis*; EB - *Eremomyces bilateralis*; VG – *Verruconis gallopava*; DA – *Dissoconium aciculare*; NA - *Neohortaea acidophila.* C) Presence-absence pattern of pPCD orthologs in four arthropods, *Folsomia candida*, *Spodoptera litura*, *Bradysia coprophila*, and *Eurytemora affinis*. Dot color and cell color indicate the feature architecture similarity to the respective seed protein using either the seed (FAS_F; dot) or the ortholog (FAS_B; cell) as reference. The number of detected co-orthologs is indicated by the green inlay.

**Table 2.**
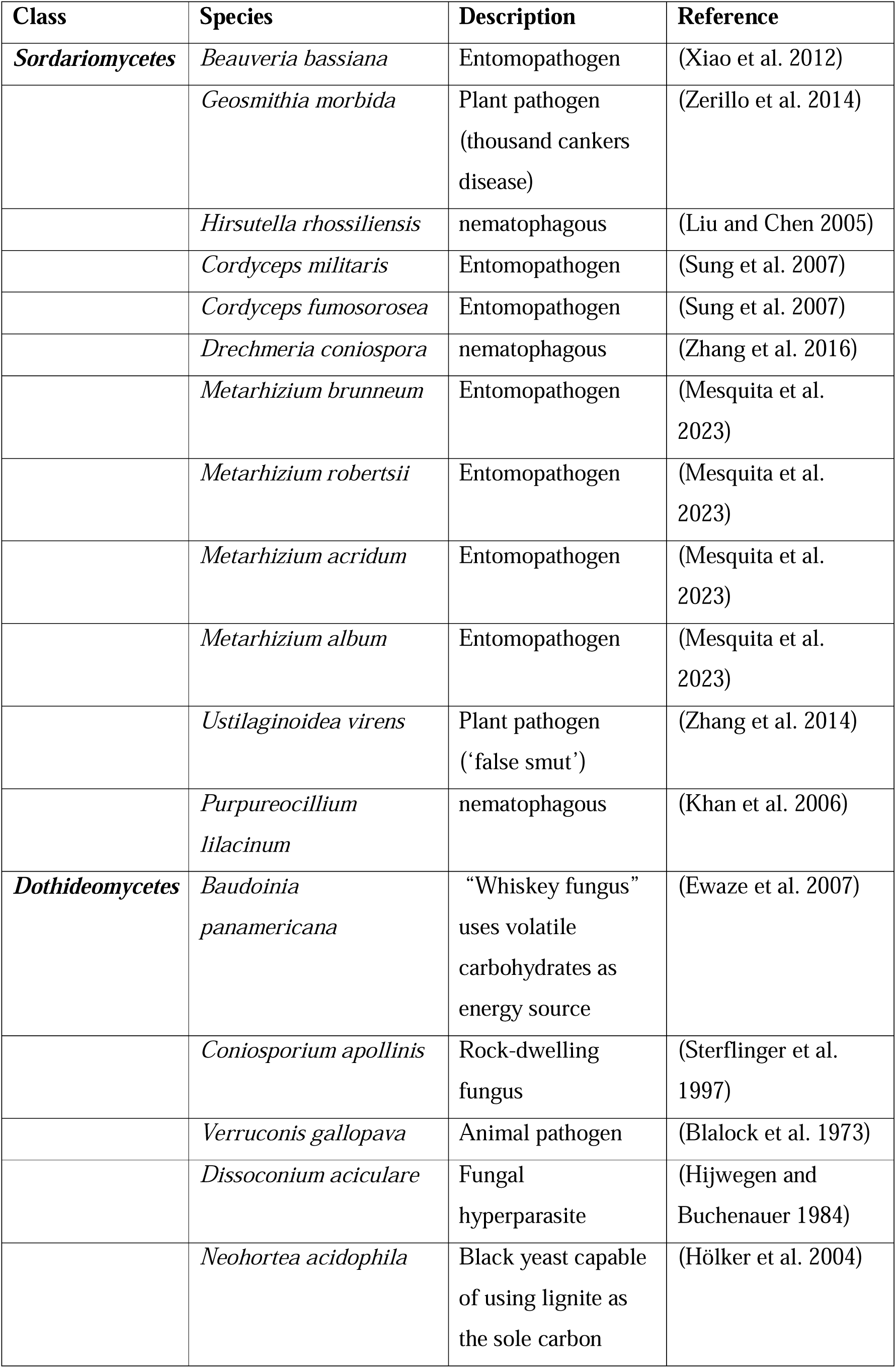
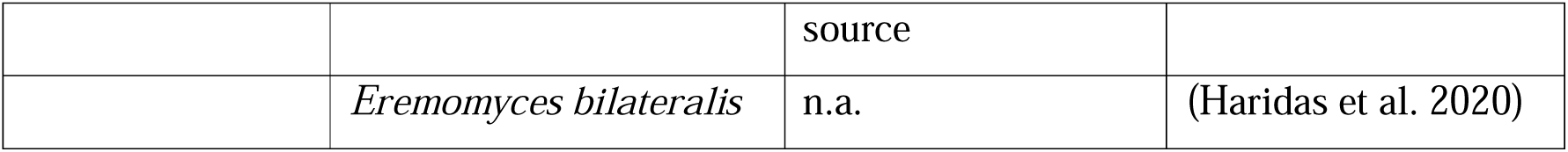
Fungal taxa with reduced numbers of pPCD orthologs

### pPCDs in animals

*Sordariomycetes* and *Dothideomycetes* are showcase examples that changes in the pPCD repertoires are readily identifiable in the UMAP and are indicative of changes in a species’ lifestyle. We next sought to find similar examples in animals. Overall, animals possess orthologs for only a few pPCDs (Fig. 3A). Those enzymes, for which orthologs are detected throughout the animals convey functions that are potentially, but not necessarily, connected to plant cell wall degradation. For example, glycosyl hydrolases of the GH1 family are often found in necrotrophic fungi where they are involved in the degradation of cellulose, hemicellulose and pectin (Chang et al. 2016). Outside fungi, we find orthologs to the *R. solani* GH1 family glycosyl hydrolase QRW24047.1 throughout the plants and the animals (Fig. S16). In plants, the detected orthologs are involved in a broad spectrum of functions including cell wall remodeling (Opassiri et al. 2006). However, their wide-spread presence in animals indicates a shift in function on this lineage. Human orthologs, for example, are annotated as lactase/phlorizin hydrolases (see Fig. S16C) and catalyze the hydrolyzation of lactate into the monomers β-D-galactose and D-glucose (Nemeth et al. 2003).

We next ranked the animals in our collection according to the diversity of their pPCD repertoires (Table S3). Among the top-ranking taxa, we find mainly arthropods and mollusks, which aligns well with reports in the literature (Pauchet et al. 2010; Tsuji et al. 2013; Busch et al. 2017; Yoshioka et al. 2017). Integrating this list with the UMAP results, four arthropod species stand out: The copepod *Eurytemora affinis*, the cotton leaf worm (*Spodoptera litura*) and the black fungus gnat (*Bradysia coprophila*)—two plant pathogens—as well as the springtail *Folsomia candida*. They are rich in pPCDs (Table S3), and they are nested in a cluster that comprises predominantly mollusks, instead of being placed in vicinity to other arthropods (Figs. 4A and B). Like the findings for the fungi, this suggests a recent shift in the composition of their pPCD repertoires. Moreover, the latter three species comprise the most diverse set of pPCDs among all investigated animals (Fig. 4C and Table S3).

### Orthologs to cell wall degrading enzymes – intrinsic genes or contaminations?

Expansions of enzyme repertoires in individual species can be indicative of evolutionarily recent gene gains. In support of this, it has been repeatedly hypothesized that animals have horizontally acquired plant cell wall degrading enzymes from either bacteria or fungi (Kirsch et al. 2014; Druzhinina et al. 2018; Busch et al. 2019). However, both fungi and bacteria are also common contaminants in genome assemblies (Steinegger and Salzberg 2020). It is conceivable that genome assemblies of animals feeding on plant material are contaminated with sequences from microbial organisms capable of cellulose degradation. Like other ortholog search tools, fDOG has no means to differentiate between genes that are in the genome of a target species or that are contaminations in the genome assembly. We therefore performed a taxonomic assignment for the pPCD orthologs in the four respective taxa. In the cotton leaf worm (*S. litura*), orthologs to 7 of the 16 represented pPCD sub-families are taxonomically assigned to bacteria. In addition, most reside on very short contigs often comprising only a single gene (Table S4). Thus, these sequences are very likely bacterial contaminations in the *S. litura* gene set. We therefore did not consider this taxon in any further analysis. The orthologs detected in the brachiopod *E. affinis* are, with one exception, assigned to arthropods (Table S5). Only the ortholog of the fungal endo-1,3(4)-beta-glucanase (GH81) appears to be of bacterial origin. Yet, we could confirm that the corresponding gene is embedded in a contig with 44 other genes and the taxonomic assignment of the flanking genes is inconspicuous (Fig. S17A). This provides evidence that this gene was recently acquired by a horizontal gene transfer. Similarly, we find no indication that contaminations account for the detection of pPCDs for the black fungus gnat (*B. coprophila;* Table S6) and the springtail (*F. candida;* Table S7), although two possible HGT events were detected for *F. candida* (PL1 and GH10; Fig. S17B-C). In summary, there is strong evidence that three out of four arthropod species have indeed significantly extended their repertoire of cell wall degrading enzymes.

### Feature architecture changes indicate functional divergence

We next increased the resolution of our analysis by investigating whether changes in the protein feature architectures provide an indication that the detected pPCD orthologs in the three arthropod species may have diverged in their enzymatic activity from the respective seed proteins. 63 out of 122 orthologs have feature architectures largely resembling that of the respective seed proteins (FAS score difference < 0.25; Table S8). The architectures of the remaining orthologs differ to an extent that changes in their activity become more likely. We will illustrate this with three examples. A major facilitator superfamily transporter of *R. solani* (UniprotID: QRW18479) is assigned to the GH51 family of glycosyl hydrolases. Next to an MFS_1 Pfam domain (PF12832), it is annotated with a C-terminal alpha-L-arabinofuranosidase Pfam domain (PF06964; Fig. 5A). Since hemi-cellulose partly contains glycosidic bonds of arabinose, QRW18479 is likely involved in its degradation. Orthologs of this protein are found in the three focal arthropods, but also throughout the eukaryotes.

**Figure 5.**
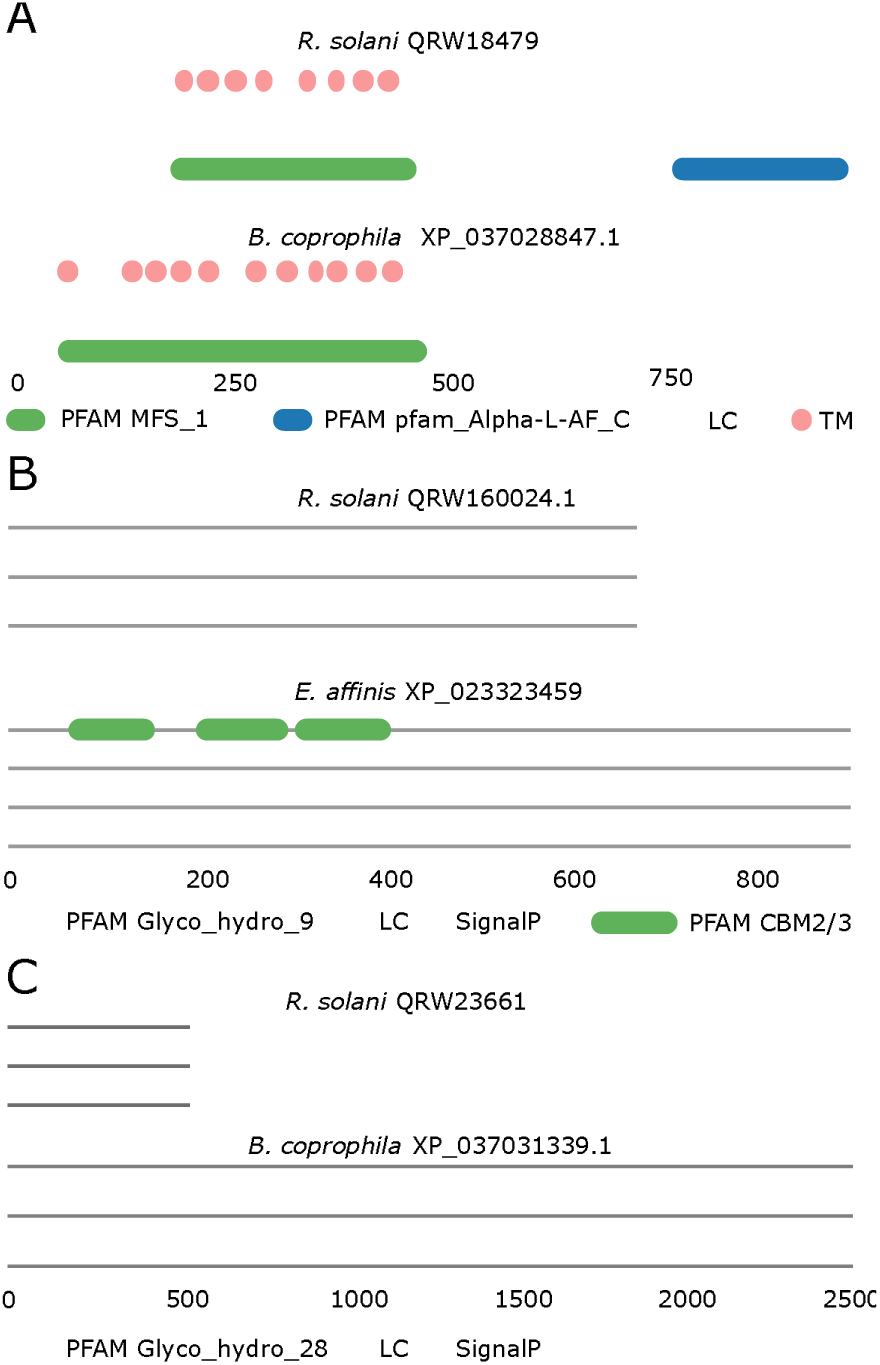
Feature architecture changes between fungal proteins and their orthologs in the arthropods suggest changes in function. A) A putative Alpha-L-arabinofuranosidase from *R. solani*. The arabinofuranosidase domain is absent from the *B. coprophila* ortholog. PFAM_MFS_1: PF07690; PFAM_Alpha-L-AF_C: PF06964. B) A putative GH9 family cellulase from *R. solani*. The ortholog in E. affinis has an additional 3 instances of a cellulose binding motif (CBM). PFAM_Glyco_hydro_9: PF00759; PFAM_CBM_2: PF00553. C) Putative endo-polygalacturonase. The *B. coprophila* ortholog has a quadruplicated domain architecture compared to the *R. solani* protein. PFAM_Glyco_hydro_28: PF00295. See protein IDs represent Uniprot Ids, whereas the IDs of the orthologs are NCBI RefSeq Accessions.

Consistently, these orthologs have low FAS_F scores indicating that a feature from the *R. solani* protein is missing (Fig. S18). A visual inspection reveals that they all lack the alpha-L-arbinofuranosidase Pfam domain. Although they may share the transporter activity with the fungal protein, they cannot exert the furanosidase activity. Taken at face value, this suggests that an evolutionarily very recent gene fusion joined the two Pfam domains on the *R. solani* lineage. Alternatively, it could also be the result of an artificial gene fusion introduced during the gene identification in the genome of *R. solani*.

The feature architecture difference observed in the *E. affinis* ortholog of a GH_9 family cellulase (UniprotID: QRW16024; FAS_B = 0.3) is likely reflecting a genuine change. The *E. affinis* ortholog shares the C-terminal GH_9 family cellulase domain and an N-terminal signal peptide with the *R. solani* protein. In addition, it harbors three cellulose binding modules (CBM_2) in its N-terminal half (Fig. 5B). This change likely does not affect the catalytic activities of the protein. However, it has been previously reported for glycoside hydrolase family 9 ‘endo-processive’ cellulases that the addition of CBMs may increase the processivity of the enzymes (Sakon et al. 1997). A similar effect is conceivable for the *E. affinis* protein. The third example is a fungal galacturonan 1,4-alpha-galacturonidase (QRW23661) which is presumably involved in pectin hydrolysis. The ortholog in *B. coprophila* is about five times larger than the corresponding seed protein from *R. solani*. It comprises all features found in the fungal protein but in multiple copies (Fig. 5C). We hypothesize that the domain multiplication in the insect protein serves to increase the dosage of the enzymatic activity. This suggests that the hydrolyzation of pectin plays a particular role for *B. coprophila*.

## Discussion

Orthology assignments integrate genes across taxa into an evolutionary context and form the basis for tracing functions across species and through time. Scope and resolution of orthology-based analyses increase with the number of taxa that can be considered in a single analysis. Given the wealth of data generated from the numerous biodiversity genomics projects worldwide, orthology assignments have become a limiting factor. Among the many tools that are available for identifying orthologs (Nichio et al. 2017; Linard et al. 2021; Nevers et al. 2022), fDOG has several unique features. Without a noticeable impact on both sensitivity and accuracy—when compared to state-of-the-art tools that require the comparison of entire proteomes—fDOG performs a profile-based ortholog search starting from a single protein sequence as a seed. Compared to other tools, such as HaMStR (Ebersberger et al. 2009), OMAmer, or EggNOG mapper, this makes it independent from any precomputed set of orthologs. By that, fDOG offers a flexibility that is, to our knowledge, unprecedented in the field of orthology prediction. At the same time, restricting the ortholog search to the subset of genes that are directly relevant for the downstream analyses reduces the energy footprint of an analysis and resembles a significant step towards Green Bioinformatics (Grealey et al. 2022). The linear runtime complexity further allows to upscale taxon sets to sizes that were thus far reserved for unidirectional homolog searches. As a second unique feature, fDOG is the only tool that integrates a feature architecture comparison into the workflow of the ortholog search.

Thus, fDOG can be used to produce taxonomically highly resolved, ortholog-based and feature architecture aware phylogenetic profiles. This facilitates to adjust the scale within a single analysis, ranging from broad-scale overviews on the kingdom and superkingdom-level summarizing the information across thousands of taxa, to detailed investigations of the presence-absence pattern of a particular feature within the ortholog of an individual strain or species. This allows to detect changes in the set of protein-coding genes that either occurred recently and are hence confined to only a few taxa, or that reflect an artefact during genome assembly or gene annotation (see (Dosch et al. 2023)). We have here demonstrated the use of fDOG in the tracing of pPCD orthologs across the tree of life. Further applications of the software together with some limitations are provided in the online supplementary material (Supplementary text 1).

### Visualization of phylogenetic profiles

Traditionally, phylogenetic profiles are displayed using taxon-gene matrices where each cell indicates whether a gene was detected in the corresponding taxon, or not (Pellegrini et al. 1999; Tran et al. 2018; Birikmen et al. 2021). In principle, this allows to directly relate the (non-)detection of a gene to its functional annotation and lastly to the individual species’ phenotypes. Summarizing phylogenetic profiles for hundreds of genes across thousands of taxa requires, however, novel approaches to visualize the resulting high-dimensional biological data. The uniform manifold approximation and projection (UMAP) (McInnes et al. 2020) is a non-linear dimension reduction technique, which has the advantage over a conventional Principle Component Analysis that it avoids distinct data clusters on overlapping areas of the graph (Becht et al. 2019). UMAP was initially introduced into biological data analysis to identify distinct cell populations in single-cell transcriptomic data. Here, we have explored the use of UMAP for generating a novel representation of phylogenetic profiles with the objective of spatially clustering taxa that share similar presence/absence patterns of the investigated genes. The results are promising as they provide an informative overview of differences and similarities in the repertoire of pPCDs across more than 18,000 taxa spanning the tree of life (see Fig. 3A). Taxonomically homogenous clusters indicate evolutionarily stable enzyme sets. In turn, changes in both size and composition of the enzyme sets inside certain systematic groups are flagged by a spatial shift of the respective taxa. One example are the Sordariomycetes, that are represented in two distinct clusters in our analysis. The different spatial localizations reflect differences in the lifestyle of the members in the two clusters (Zhang et al. 2014). Consequently, the UMAP visualization of phylogenetic profiles allows the user to identify individual species of interest that warrant more refined analysis, even in analyses that span thousands of taxa.

### Catalogues of plant cell wall degrading enzymes

The degradation of primary plant cell walls requires a diverse cocktail of enzymes, comprising cellulases, hemi-cellulases and pectinases. Many organisms equipped with these enzymes decompose dead plant material (Keggi and Doran-Peterson 2020; Griffiths et al. 2021). They make the stored energy accessible and contribute to releasing the bound carbon, thus contributing to the global carbon cycle. Some are also potent plant pathogens with substantial impacts on crop yield and therefore are of considerable economic importance.

Irrespective of their ecological impact, all are relevant inputs for biotechnological applications that seek to find solutions to utilize plant material as a source of many different compounds (Benedetti et al. 2019; Giovannoni et al. 2020; Matrose et al. 2021). The CAZy database (Drula et al. 2022) provides an extensive catalogue of carbohydrate active enzymes across in a diverse collection of species. We have used this data as a basis to compile the most comprehensive taxonomic overview of enzymes that can contribute to primary plant cell wall degradation, to date. Compared to the CAZy database (Drula et al. 2022), we see three advantages. First, we focus on taxa with annotated genome sequences available. This provides a more complete and systematic view on a species’ enzyme repertoire because the entire gene set can be investigated. Consequently, the capabilities of an organism to degrade plant cell walls can be better assessed. For example, as of October 2024, *F. candida* is represented with only two sequences in the CAZy database, both representing GT3 family glycosyl transferases. This is at odds with previous findings providing strong evidence for a functional GH45 family cellulase in this organism (Muelbaier et al. 2024), and with the results of this study where we provide evidence that *F. candida* is equipped with a rich set of pPCDs (see Fig. 4). Second, the phylogenetic profiles reveal gene duplications and losses, but also the unexpected presence of orthologs in taxa that is hard to explain by a vertical inheritance of the corresponding genes. Such instances flag cases where further curation is necessary to differentiate between gene acquisition via horizontal gene transfer, and the presence of contaminations in a genome assembly. Third, the integration of orthology assignments with feature architecture comparisons provides a solid basis to curate the distribution of patterns in a semi-automated manner. This allows to identify cases where the presence of an ortholog most likely does not coincide with the presence of its function (see Fig. 5).

### Plant cell wall degrading enzymes in animals

Gradually, evidence emerges that a small and taxonomically diverse set of invertebrates possess intrinsic cellulase-encoding genes in their genomes (Gilbert 2010). Many of the hitherto described cellulases are members of the GH45 family (Girard and Jouanin 1999; Song et al. 2017; Busch et al. 2019). In-depth analysis of individual taxonomic groups, such as the polyphaga, the horn mites (Oribatida) and the springtails (Collembola) have furthermore revealed that GH45 family cellulases are widespread within these groups (Busch et al. 2019; Muelbaier et al. 2024). This indicates an evolutionarily early acquisition of the corresponding genes, probably by a horizontal transfer from fungi (Busch et al. 2019; Muelbaier et al. 2024). Although GH45 family enzymes with their endo-β-1,4-glucanase activity can degrade various polysaccharides, such cellulose, lichenin and xyloglucan (Busch et al. 2019), their activity alone is not sufficient to degrade primary plant cell walls. More comprehensive characterizations of plan cell wall degrading enzymes have, thus far, been largely restricted to the insects (Pauchet et al. 2010; Calderón-Cortés et al. 2012; McKenna et al. 2016). This has revealed that polyphaga, a suborder within beetles, possess a considerably rich spectrum of cellulases and pectinates (Pauchet et al. 2010). Our results are the first step towards comprehensively charting the presence of enzymes potentially involved in plant cell wall degradation across animals. This confirm that vertebrates are devoid of intrinsic cellulases. We find that several taxa, such as hemichordates, echinoderms, crustaceans, mollusks and cnidaria are equipped with various combinations of pPCDs (Table S3). The black fungus gnat, *Bradysia coprophila* (Sciaridae, Diptera), however comprises the richest and most diverse collection of intrinsic pPCDs across all animals analysed by us. This finding aligns well with a recent report in which manual curation revealed that *Pseudolycoriella hygida*, another member of the Sciaridae, is particularly rich in plant cell wall degrading enzymes (Trinca et al. 2023). Many of these enzymes are represented in the saliva of the *P. hygida* larvae, which feeds on plant litter, and it was shown that this enzyme mix contributes to the degradation of the plant material. Of the 27 pPCD sub-families identified by our ortholog search, we find that 17 are consistently identified by the study of Trinca et al. (2023). For 7 of these, proteomic evidence exists that they are indeed expressed in the larval saliva (Trinca et al. 2023). This demonstrates the power of fDOG in tracing proteins and their function. However, the resolution of the fDOG analysis extends beyond the identification of members of an enzyme family. For example, while Trinca et al. (2023) only stated the presence of a GH28 family pectinase in black fungus gnats, our results reveal that these proteins are orthologous to the GH28 family protein in *R. solani* (Uniprot ID: QRW23661).

Compared to the fungal protein, however, the fly protein has a very specific architecture where the features of the fungal protein have been quadruplicated (see Fig. 5C), an architecture that is unique among all GH28 family members included in our study. This information is essential for investigating the precise role of this enzyme in *B. coprophila*. But it also helps in the identification of promising novel enzymes for the use in biotechnological applications.

The wide-spread presence of GH45 family cellulases in springtails (Collembola) has recently triggered speculations that springtails could enzymatically degrade plant litter (Muelbaier et al. 2024). Here we could show that *F. candida* is equipped with orthologs to the second most diverse set of plant cell wall degrading enzymes observed in the animals, thus far (see Fig. 4C). This aligns well with the large and diverse food spectrum of Collembola, including litter, pollen, algae, leaves, roots (Potapov et al. 2022). Collembola might directly feed on polysaccharides deposited in the plant cell walls, or their cellulases contribute to the enzymatic degradation of cell wall polysaccharides opening access to the cytosol inside.

Another and presumably more important food source might be fungi that colonize the inside of plant cells in plant litter. Feeding on these fungi would require at least a partial digestion of the surrounding plant material.

Soils contain large amounts of plant litter as part of soil organic matter, resulting in soils being the major sink of terrestrial carbon (Jones and Donnelly 2004). Decomposition of plant material thus represents an important driver of global carbon cycling. Therefore, the three taxa highlighted in this study should be considered as players in of global carbon cycling that have been hitherto overlooked.

## Materials & Methods

### Benchmark Data

For benchmarking we used the Quest for Orthologs (QfO) reference datasets 2020_04 which were obtained from the EMBL-EBI FTP server (http://ftp.ebi.ac.uk/pub/databases/reference_proteomes/). The pairwise orthology assignments between human and other QfO taxa were filtered from the benchmark release 2020 downloaded from the OrthoBench websites (https://orthology.benchmarkservice.org/). These 17 methods were taken into account, including RBH/BBH, Hieranoid 2, InParanoid, MetaPhOrs v.2.5, OMA Groups, OMA Pairs, OrthoFinder 2.5.2, OrthoInspector 3, OrthoMCL, PANTHER 16.0 all, PANTHER 16.0 LDO, PhylomeDB V5, RSD, SonicParanoid with 4 different settings default, fast, most sensitive and sensitive. From the publicly available data, we selected for the benchmark the subset of ortholog pairs that include a human protein.

### pPCD data

Gene sets for the search for pPCD orthologs were downloaded from the RefSeq section of the NCBI genome database. The taxon collection comprised 18,565 taxa representing 16,324 bacteria, 887 archaea, and 1354 eukaryota (138 plants, 416 fungi, 429 vertebrates and 262 invertebrates and 109 other taxa).

### Protein sequence comparison

Feature architecture similarities are computed with FAS (Dosch et al. 2023) using the default feature set comprising PFAM and SMART domains (Letunic et al. 2021; Mistry et al. 2021), transmembrane helices predicted with TMHMM 2.0c (Krogh et al. 2001), signal peptides with SignalP 4.1 (Petersen et al. 2011), coiled-coils structures with COILS2 (Lupas 1996), low complexity regions with SEG (Wootton and Federhen 1993) and fLPS 2.0 (Harrison 2021).

### Detecting gene duplications on the fungal lineage

For each paralogous group of genes for which a fungal organism served as seed species, we traversed the taxonomic tree of target species from the seed species to its most distantly related clade using the ete3 toolkit (Huerta-Cepas et al. 2016). We approximate the origin of the gene duplication by identifying the closest relative of the seed species for which the same gene is found as ortholog to two different seed genes in the paralogous group. If the gene duplication occurred in the Fungi lineage, the seed gene with the most orthologs in the Metazoan lineage was kept as representative.

### Tree topology testing

Multiple sequence alignments were performed with MUSCLE 5.1 (Edgar 2022) using default options. Phylogenetic trees were computed using IQ-TREE v2.2.0 (Minh et al. 2020) allowing the algorithm to automatically select the best-fitting substitution model with ModelFinder (Kalyaanamoorthy et al. 2017). Testing of competing phylogenetic hypotheses were performed using the AU test (Shimodaira 2002) as implemented in IQ-TREE at a significance level of 5%. We corrected for multiple testing using the False Discovery Rate (Benjamini and Hochberg 1995).

### Hardware and parameter settings

Computations was run using a computer node with 2 processors AMP EPYC 7601, each containing 32 cores operating at 2.2Ghz, and 1TB of RAM. Unless otherwise noted, we ran fDOG with the set of default parameters.

### Visualization

Phylogenetic profiles of orthologs detected with fDOG were visualized and analyzed using PhyloProfile v1.18 (Tran et al. 2018). Orthologs with a FAS-score below 0.3 were filtered out and the remaining phylogenetic profiles were transformed into binary presence/absence vectors (1 if at least one ortholog was found, otherwise 0). These vectors were subsequently projected into 2D space using the UMAP algorithm (McInnes et al. 2020).

### Detection of contaminating genes

Each gene in a gene set of a give species was taxonomically assigned with DIAMOND v2.1.9 (Buchfink et al. 2021) using the NCBI non-redundant protein database (downloaded on 17^th^ of July 2023) as a reference. If multiple isoforms were annotated for a gene, we used the longest isoform as a query in the DIAMOND search. The hit of the query sequence against itself was excluded from the hit list, and the bit score margin to include lower ranking hits into the taxonomic assignment was kept at the default value of 10% of the best hit’s bit score. To put the taxonomic assignment into a context of the corresponding genome assembly, we assigned each gene to the contig it was annotated on. A gene was tentatively annotated as ‘foreign’ if it was assigned to a taxon that is not on the path from the species the genome assembly was derived from to the root of cellular organisms. A foreign gene was considered a contamination (i) if the length of the scaffold it resides on is smaller than the mean scaffold length across the assembly and (ii) if its direct neighbors were also flagged as ‘foreign’. Otherwise, it was tentatively flagged as ‘horizontally acquired’.

## Notes

https://applbio.biologie.uni-frankfurt.de/download/fDOG_pPCD/

### Competing Interest Statement

The authors have declared no competing interest.

### Summary of Updates

We have added a comprehensive workflow describing the way how fDOG works as a new supplementary figure 1.

