## Supplementary information for "Tracing the taxonomic distribution of plant cell wall degrading enzymes across the tree of life using feature architecture aware orthology assignments"

### Online Supplementary Information

#### Supplementary text

##### fDOG – further applications

fDOG is a versatile method to establish feature architecture-aware phylogenetic profiles using orthology assignments for custom-tailored gene- and taxon sets. Here, we have demonstrated the application of fDOG in the tracing of plant cell wall-degrading enzymes across the tree of life. However, applying the same approach to any other set of functionally interacting proteins is straightforward. Additionally, fDOG can be used for any other application that is based on the generation and interpretation of phylogenetic profiles. For example, any collection of phylogenetically informative marker genes can serve as input for fDOG, rendering the compilation of a data matrix for phylogenomic analyses a routine task. Keeping track of the number of co-orthologs identified in the individual taxa makes it straightforward to identify and cope with lineage-specific gene duplications that may render the interpretation of the phylogenetic signal problematic. Likewise, fDOG can be used in a BUSCO-like fashion to determine the presence-absence pattern of core genes in a phylogenetic clade. Outliers in the FAS score can then indicate cases where the gene prediction likely resulted in spurious gene models (see Dosch et al. 2023). From the results, gene set completeness estimates across the subsumed taxa can be readily computed. In addition, the visualization of the resulting phylogenetic profiles allows an intuitive interpretation of the results, where e.g., missing or additional features in the identified orthologs can aid in the identification of errors in the gene prediction. This is especially facilitated by the seamless integration of fDOG with PhyloProfile (Tran et al. 2018).

##### Limitations of fDOG

fDOG identifies orthologs to a seed protein with sensitivities and specificities comparable to state-of-the-art ortholog search tools that require entire proteomes as input. In addition, the scoring of the feature architecture similarities between orthologs as a proxy of their functional diversification is, to our knowledge, a feature that is unique to fDOG. However, fDOG brings along also some limitations. As it is the case for any software that identifies orthologs between pairs of species, the orthology assignments of fDOG hold only between the seed protein and its orthologs in one target species. Other than that, fDOG makes no inferences about the precise evolutionary relationships of the proteins subsumed in a phylogenetic profile. fDOG further considers individual seed proteins as separate evolutionary entities. Therefore, it is possible that paralogous seed proteins share, in parts, the same orthologs in those target taxa that diversified prior to the gene duplication event that gave rise to the paralogs. In our example application, this is seen by the 236 pPCDs from the six seed taxa, which reduce to only 76 distinct evolutionary lineages in the last common ancestor of the fungi. Eventually, feature architecture differences are only a proxy for the functional diversification of orthologs, because it is largely unexplored how the differential presence, e.g., of low complexity regions, affect protein function on a global scale (see Dosch *et al.*, 2023 for a more detailed discussion). In this context, manual curation is still essential, putting a strong focus on the visual interpretation of the phylogenetic profiles.

#### Supplementary Figures

Figure S1. Targeted and feature architecture aware ortholog search with fDOG **(A)** Workflow of the profile-based ortholog search to extend a core-ortholog group C with sequences from a query species. The workflow for compiling the set of core orthologs C, and the corresponding profile hidden Markov model (pHMM_C_) that serve as input for the ortholog search is shown in (B). In brief, a pHMM based search is performed to identify significantly similar sequences to the entries in C in a query species. The resulting hit list (o_1_,…,o_n_) is ordered according to decreasing domain bit scores (bit). The hit list is cropped retaining only candidates with a domain bit score >= α x bit(o_1_). Each candidates o_1_,…,o_k_ is then used as query for a sequence similartiy based search in the protein set of the seed species (see (B)). The current implementation of fDOG uses BlastP (Altschul et al. 1990). The query protein is assigned as an ortholog to c_0_ if either of the two conditions are met: (i) the best hit of the BlastP search is the seed sequence c_0_ or (ii) the Kimura distance (Kimura 1984) between the best hit of the BlastP search and the seed sequence c_0_ is smaller than the Kimura distance between the query and the best blast hit. If neither of the two conditions is met, the query is discarded. The fDOG ortholog search concludes if all candidates for a species have been evaluated. The subsequent evaluation of the pairwise feature architecture similarities between the seed protein c_0_ and all accepted orthologs (Dosch et al. 2023) is not shown. **(B)** Workflow for iteratively compiling a core ortholog group C. The algorithm takes as input a single amino acid sequence c_0_ as seed sequence, the corresponding seed species p_0_, a set of core taxa, and a minimal and maximal taxonomic distance to control the taxonomic diversity of the core ortholog set. The set of core taxa T is cropped by removing taxa whose taxonomic distance to p_0_ is either below the minimal or above the maximal distance specified by the user. The protein set of p_0_ is extracted and it is used to create a BlastP database (see (A)). fDOG accepts c_0_ only then as a seed if either of the two conditions are met: (i) The sequence id of c_0_ is represented in the protein set of p_0_, or (ii) the best BlastP hit in the protein set of p_0_ using c_0_ as query has a length difference of no more than 10 amino acids. A BlastP e-value cut-off of 0.001 is applied. The core ortholog group C is initalized with c_0_ and the set of primer taxa P is initalized with p_0_. The candidate list is initialized as an empty set, and the current best candidate score (candScore) is initialized with 0. Next, a profile based ortholog search (see (A)) is performed in each taxon within T, where taxa t_i_ are visited in increasing taxonomic distance from p_0_. If no ortholog is detected in t_i_, the next taxon t_i+1_ is searched. Otherwise, we compute a normalized pairwise alignment score AS_norm_(c_0_,c_i_) = (AS(c_0_,c_i_)/AS(c_0_,c_0_)) where AS(c_0_,c_i_) is the pairwise alignment score of the two amino acid sequences c_0_ and c_i_ and AS(c_0_,c_0_) is the maximally possible alignment score. We compute the feature architecture similarity FAS(c_0_,c_i_) only if AS_norm_(c_0_,c_i_) x candScoreFactor is larger than the currently best candScore - 1. This reduces the computational overhead for c_i_, which cannot surpass the current best candScore even when their feature architecture is identical to that of c_0_ (FAS(c_0_,c_i_)=1). The candScore of c_i_ is computed as AS_norm_(c_0_,c_i_)+FAS(c_0_,c_i_), and c_i_ together with candScore_ci_ is added to the candidate list. The candScores are subsequently used to rank the core ortholog candidates in the candidate list. If candScore_ci_ > current best candScore, the latter variable is updated. We stop the iteration and directly append c_i_ to C and t_i_ to P if candScore_ci_ * candScoreFactor ≥ 2. The candScoreFactor is set to 1.05 per default, and can be adjusted by the user. Otherwise, we traverse all taxa in T, and the highest scoring entry in the candidate list is appended to C and the corresponding taxon is appended to P. In case of a draw, the taxonomically closer sequence to c_0_ is chosen. The sequences in C are aligned, a new pHMM is trained and the search for the next core ortholog starts, ignoring all taxa in T that are taxonomically closer to P_0_ than the taxon p_n-1_ that was appended to P in the previous iteration. The core set compilation ends if (i) the user-specified number of core orthologs has been found, or (ii) the set of candidate core orthologs is empty.

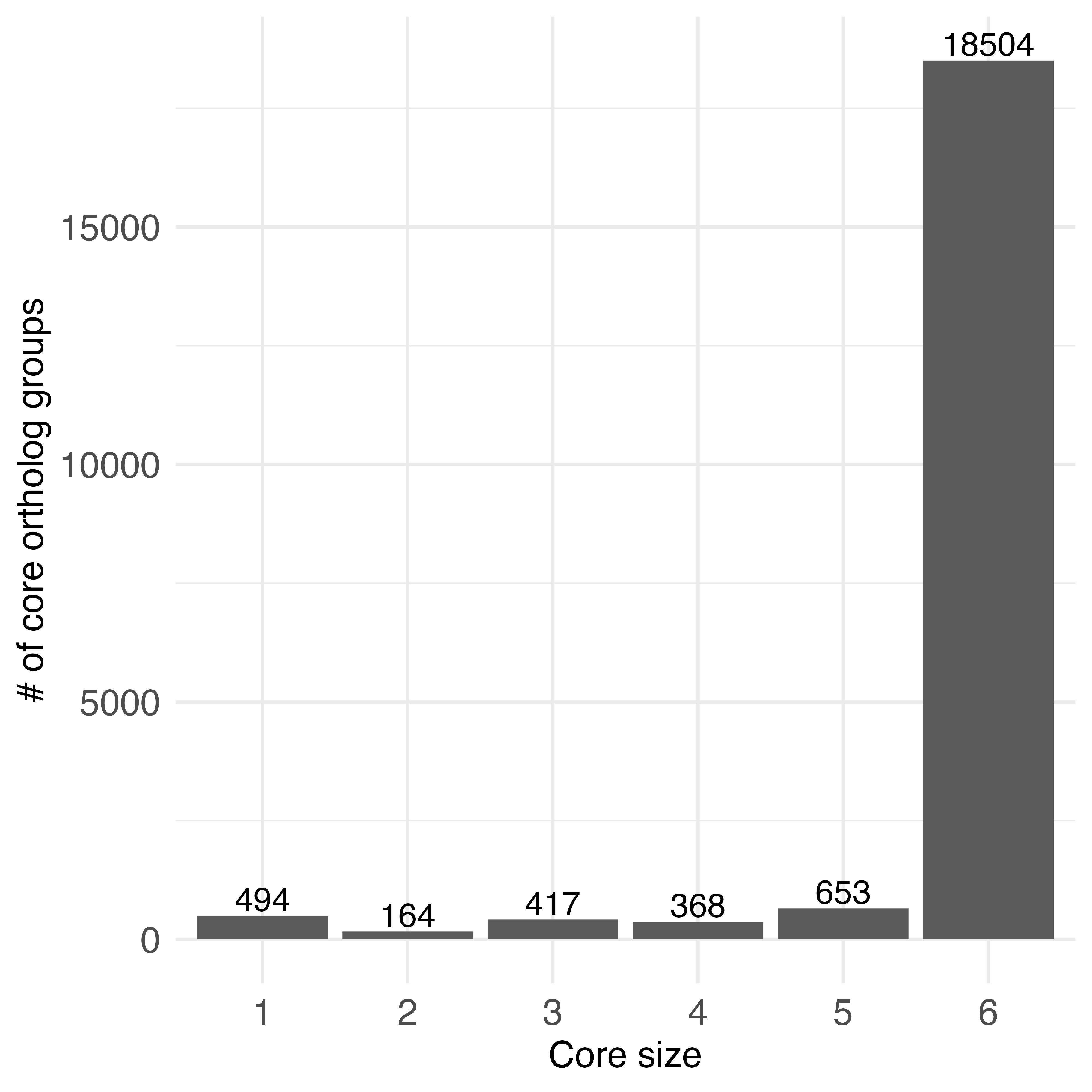

Figure S2. Group size of core ortholog sets for 20,600 human seed genes. Each core ortholog group contains up to five orthologous sequences and additionally the seed gene. The size of the core group is defined by the user (default 6). Single sequence core ortholog groups contain the human seed sequence only.

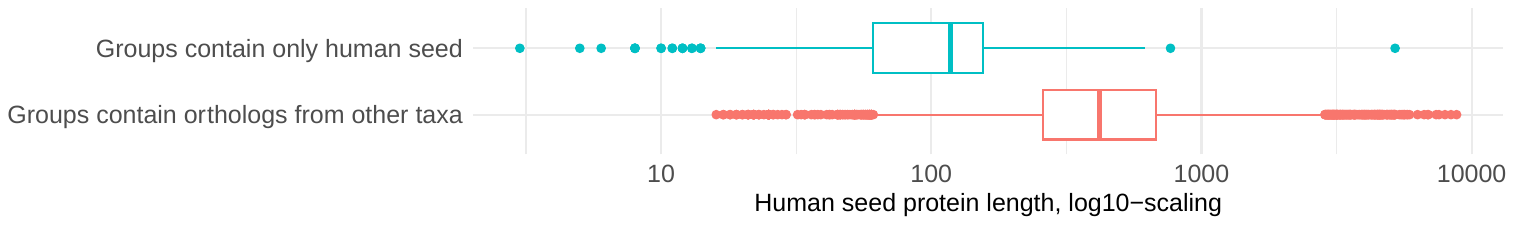

Figure S3. Human seed protein length comparison between core orthologous groups of different taxonomic diversity. The box plot in cyan represents the seed protein length distribution for 494 core ortholog groups that contain only the human sequence. The red box plot represents seed protein lengths for core ortholog groups with at least one protein from an additional species. Two proteins (Q8WXI7 with 14,508 amino acids and Q8WZ42 with 34,351 amino acids) have been excluded from the distribution plot. The mean length of the single core groups (130 amino acids) is significantly shorter than the one of the multiple core groups (564 amino acids) with p-value < 2.2e-16 (Welch’s t-test)

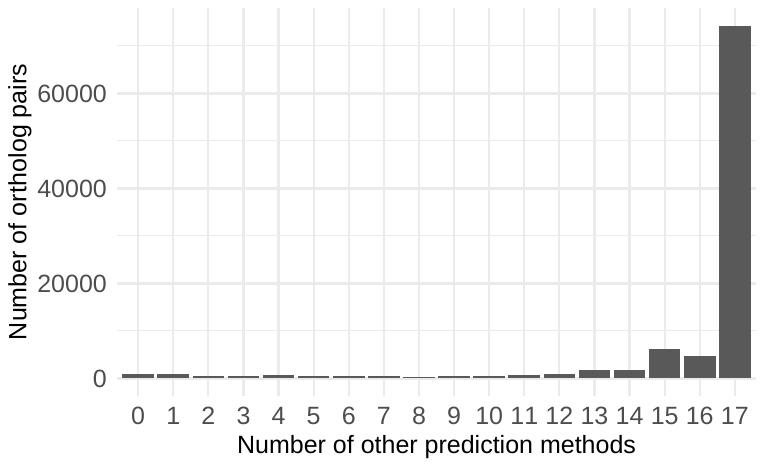

Figure S4. Consistency of orthology assignments of fDOG and other 17 orthology prediction methods. Only 1% (982 out of 97,234) of the ortholog pairs are exclusively predicted by fDOG.

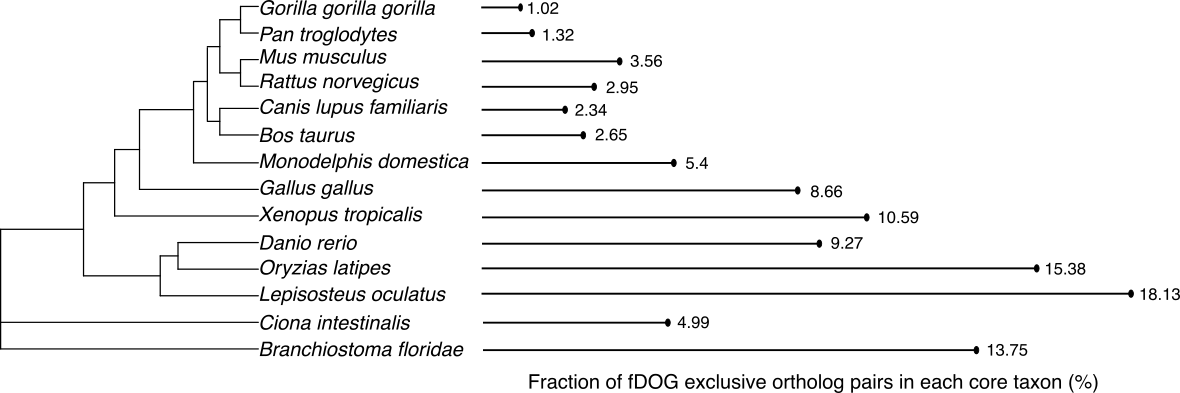

##### *Figure S5. Fraction of fDOG exclusive ortholog pairs in each core taxon in each species, sorted by the increasing taxonomic distance to human.*

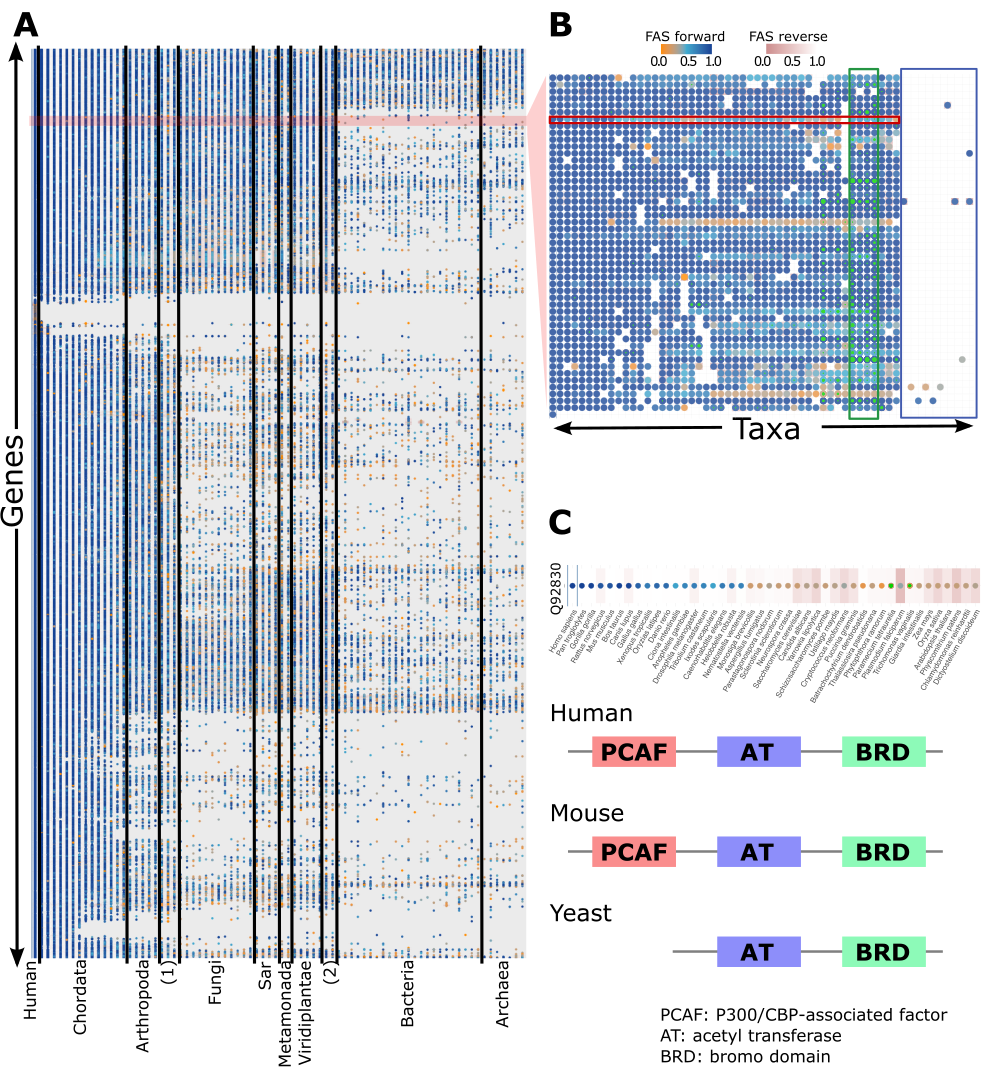

Figure S6. Phylogenetic profiles of 20,600 human proteins across 77 QfO reference taxa. **(A)** shows the full set, **(B)** a subset of 50 genes. Genes are represented in the rows and taxa in the columns ((1) – Helobdella, Nematostella, Monosiga; (2) – Dictyostelium, Leishmania). A dot indicates that an ortholog to the human seed protein was found in the respective species. Dot and cell color inform about the feature architecture similarity of the ortholog using in turns the seed protein (dot color) and the ortholog (cell color) as reference. Co-orthologs are denoted by a green inner circle. Within the green box in (B), numerous green dots can be observed, which likely echo whole genome duplications in plants. **(C)** Phylogenetic profile of the human protein Q92830, together with the feature architectures of the human, mouse, and yeast orthologs. This example illustrates the evolutionary history of the mammalian PCAF domain in the histone acetyltransferase complex (Baker and Grant 2007; Nagy and Tora 2007; Ononye and Downey 2022)

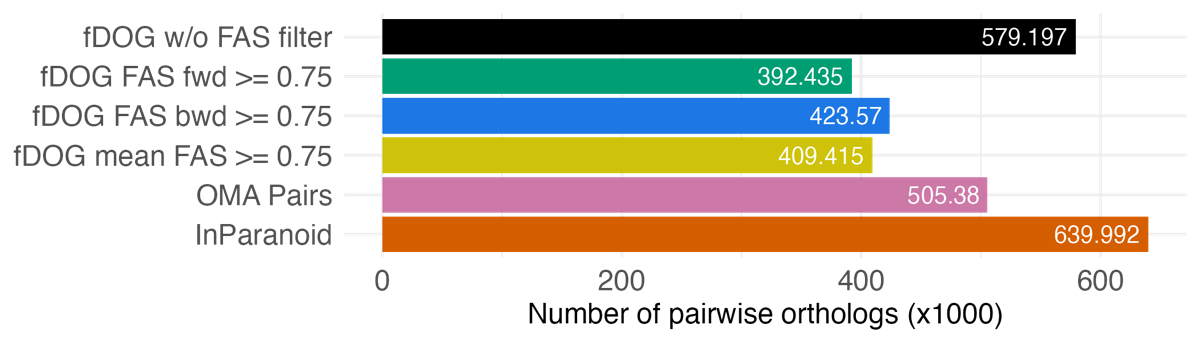

##### *Figure S7. Number of pairwise orthologs between human and the 77 other taxa in the QfO reference proteome collection for different ortholog predictors*

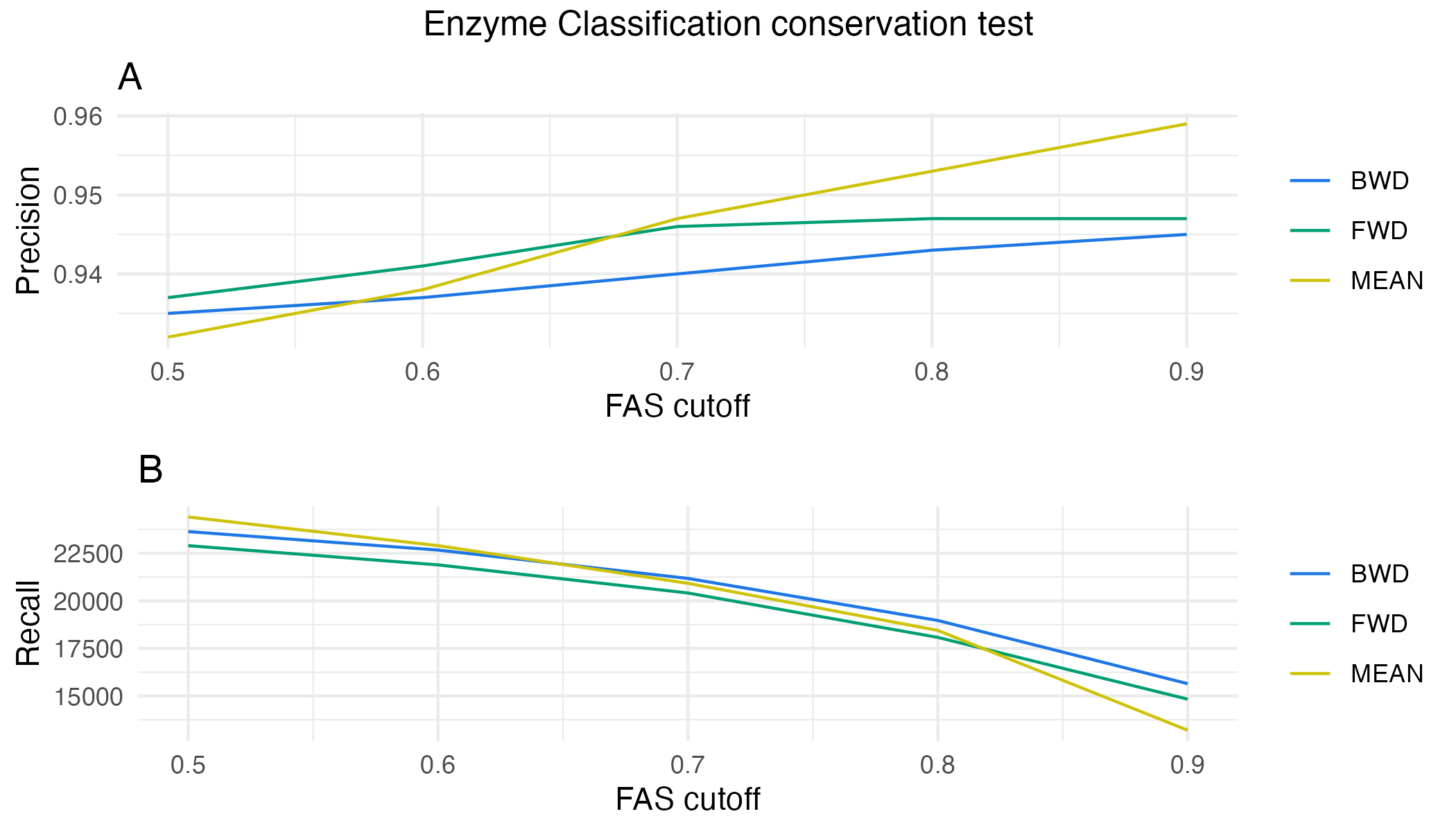

Figure S8. The effect of using different FAS filter modi and FAS cutoffs on the Enzyme classification conservation benchmark. **(A)** Precision; **(B)** recall. BWD – FAS_B computed using the ortholog feature architecture as reference; FWD – FAS_F computed using the seed protein feature architecture as reference; MEAN – (FAS_F + FAS_B)/2

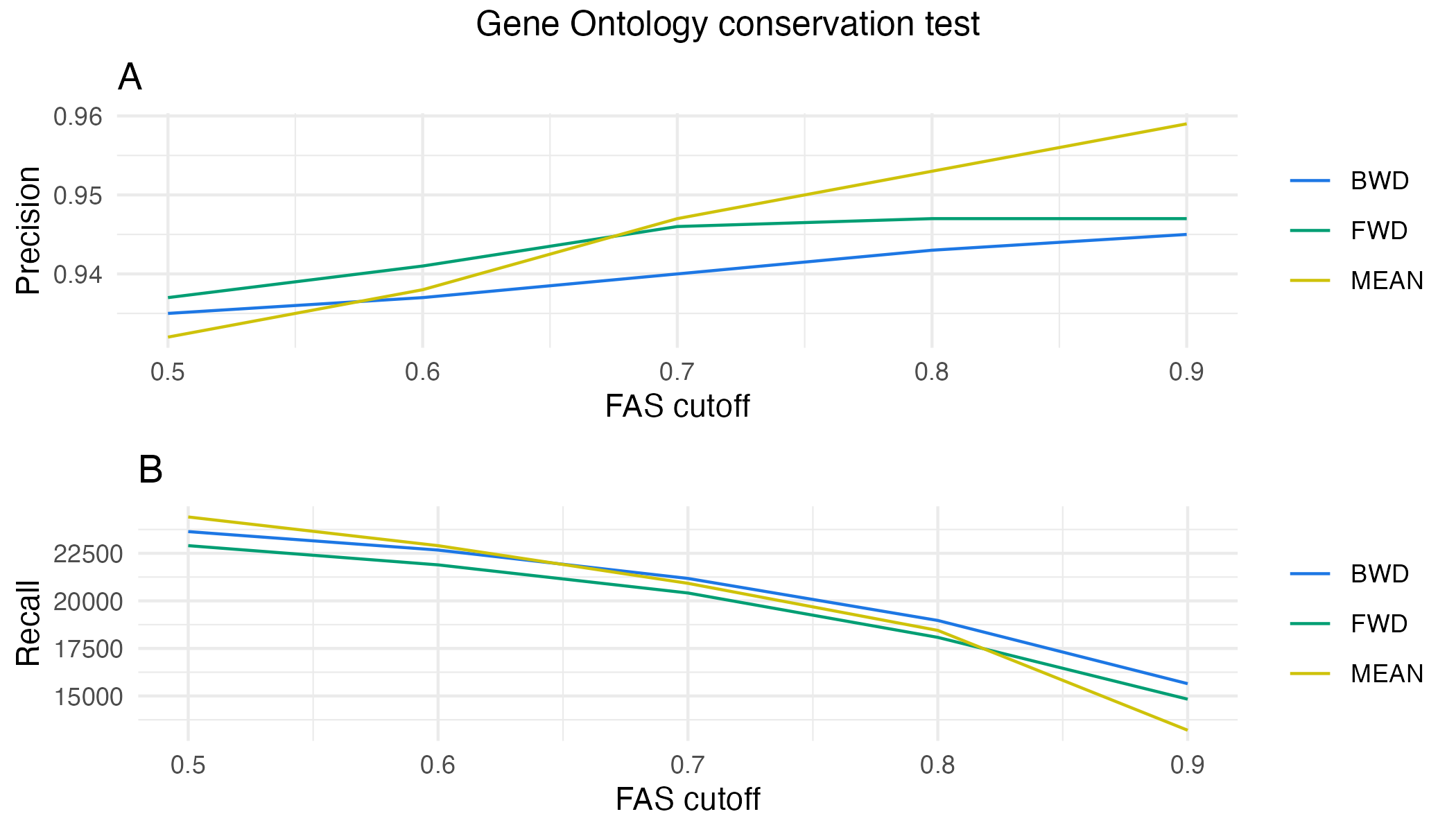

Figure S9. The effect of using different FAS filter modi and FAS cutoffs on the Gene ontology conservation benchmark. **(A)** Precision; **(B)** recall. BWD – FAS_B computed using the ortholog feature architecture as reference; FWD – FAS_F computed using the seed protein feature architecture as reference; MEAN – (FAS_F + FAS_B)/2

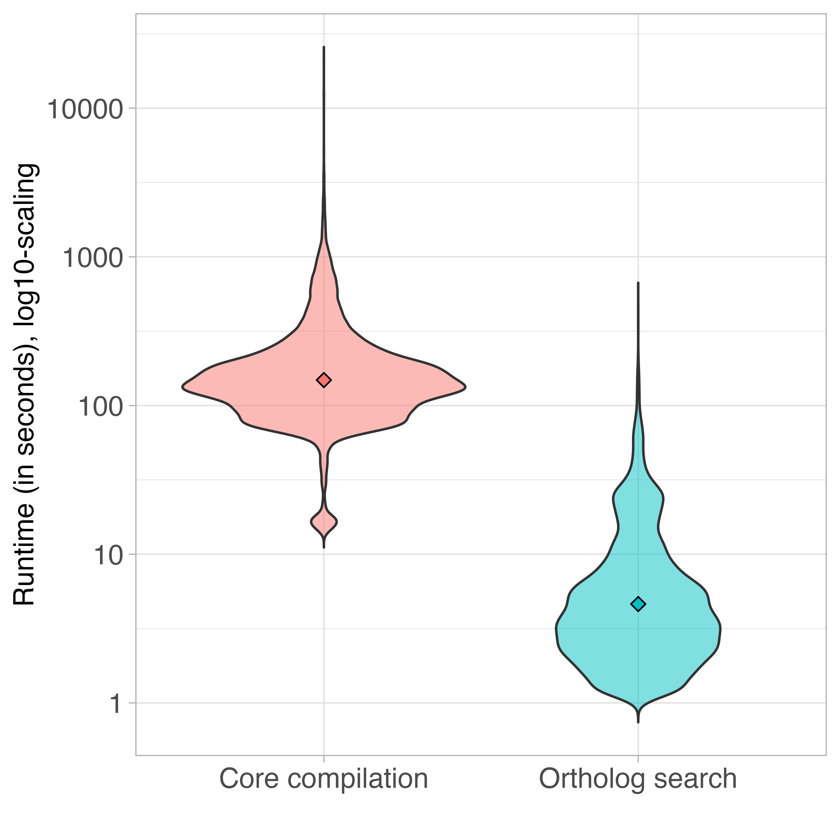

Figure S10. Run time comparison of core ortholog compilation and full ortholog search in fDOG. Log of runtime (in seconds) of the core compilation with (red) or without (green) FAS filtering and ortholog search (blue) step in fDOG. For 20,599 human proteins, the core compilation required from 13s (A0A0A0MTA1, 12 amino acids) to 3563s (A6NN14, 1253 amino acids), while the ortholog search needed only between 1s (P0CJ74, 25 amino acids) and 482s (A0A0B4J234, 113 amino acids) to be finished. On average, each sequence needed 183s for the core compilation and 8s for the ortholog search in all 78 QfO reference taxa.

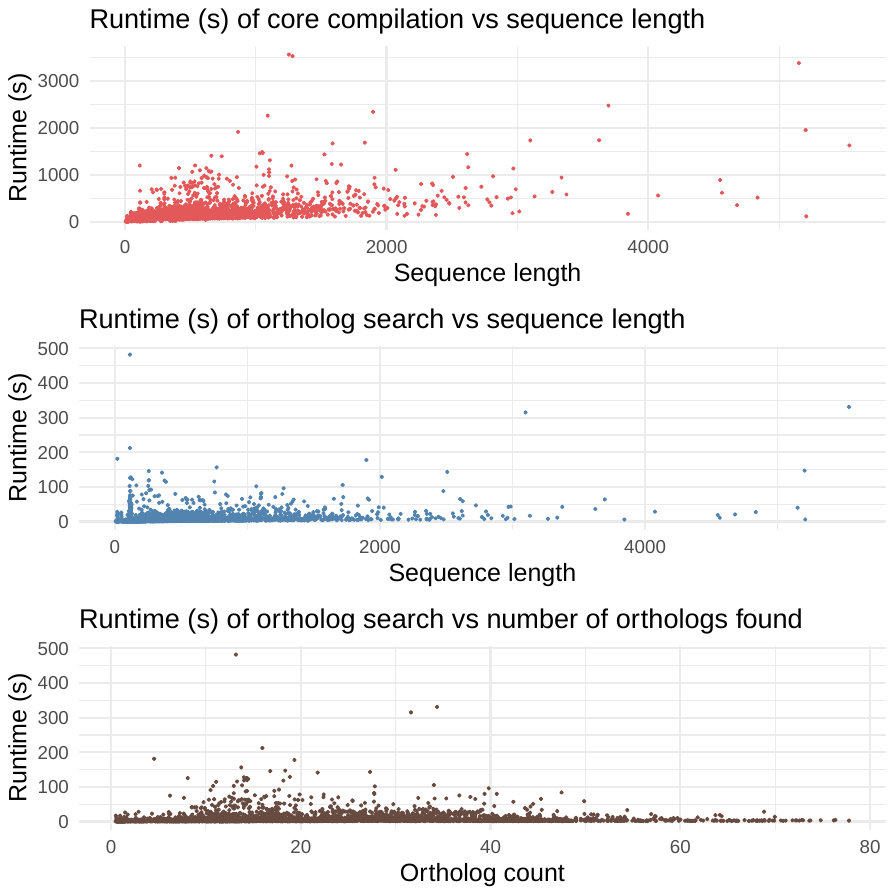

Figure S11. Parameters affecting the runtime of fDOG. Runtime (seconds) of the core compilation against the seed sequence length (red), and runtime (seconds) of the ortholog search against the seed sequence length (blue) and the number of orthologs found (brown). Only the runtime of the core compilation shows a moderate correlation with the sequence length (Pearson p-value < 2.2e-16 and correlation coefficient of 0.52). The runtime of the ortholog search has a negligible correlation with the sequence length (Pearson p-value < 2.2e-16, correlation coefficient of 0.26) and no correlation with the number of orthologs found (Pearson p-value 0.058, correlation coefficient of 0.02)

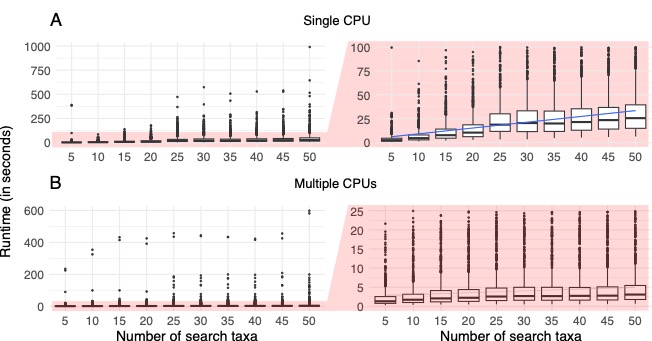

Figure S12. Runtime scaling of the fDOG ortholog search. The histograms represent the distribution of runtimes over 1000 human proteins in taxon sets varying in size from 5 to 50 taxa. **(A)** Ortholog search using a single CPU. The runtime of ortholog searches in fDOG scales approximately linearly with the number of search taxa. The left plot shows the full runtime range. To give a better resolution, the plot to the right truncates the data to consider only runs having completed after 100 seconds. **(B)** The plots represent the same analysis as in (A), but this time fDOG was run in parallel assigning the search in each taxon its own CPU. This leaves the runtime by and large invariant across different numbers of taxa.

*Figure S13. Phylogenetic profiles of the representative pPCDs across 18,000+ taxa displayed in a conventional taxon-gene matrix.* *Rows represent the 76 pPCDs, columns the individual taxa. A dot indicates the detection of an ortholog to a pPCD in the respective taxon, where the dot color informs about the extent of feature architecture similarity from a maximum of 1 (blue) to a minimum of 0 (orange).*

*Figure S14. Interactive UMAP plot of the taxa investigated in this study. The spatial arrangement of the 18,565 taxa is determined by the presence of orthologs to the 76 representative pPCDs. Only orthologs with a FAS score of at least 0.7 are considered. Dot diameter is proportional to the number of pPCD seed proteins represented by at least one ortholog. A single click on the taxa on the legend toggles their display in the UMAP. A double click on a taxon label in the legend hides all other taxa from the UMAP. Hovering over a dot in the UMAP provides information about the taxonomic label, the species name together with its NCBI taxonomy id, the number of pPCD seed proteins represented by at least one ortholog, as well as a list of the corresponding seed proteins.*

*See file file Fig-S14-UMAP-all_76-03.html*

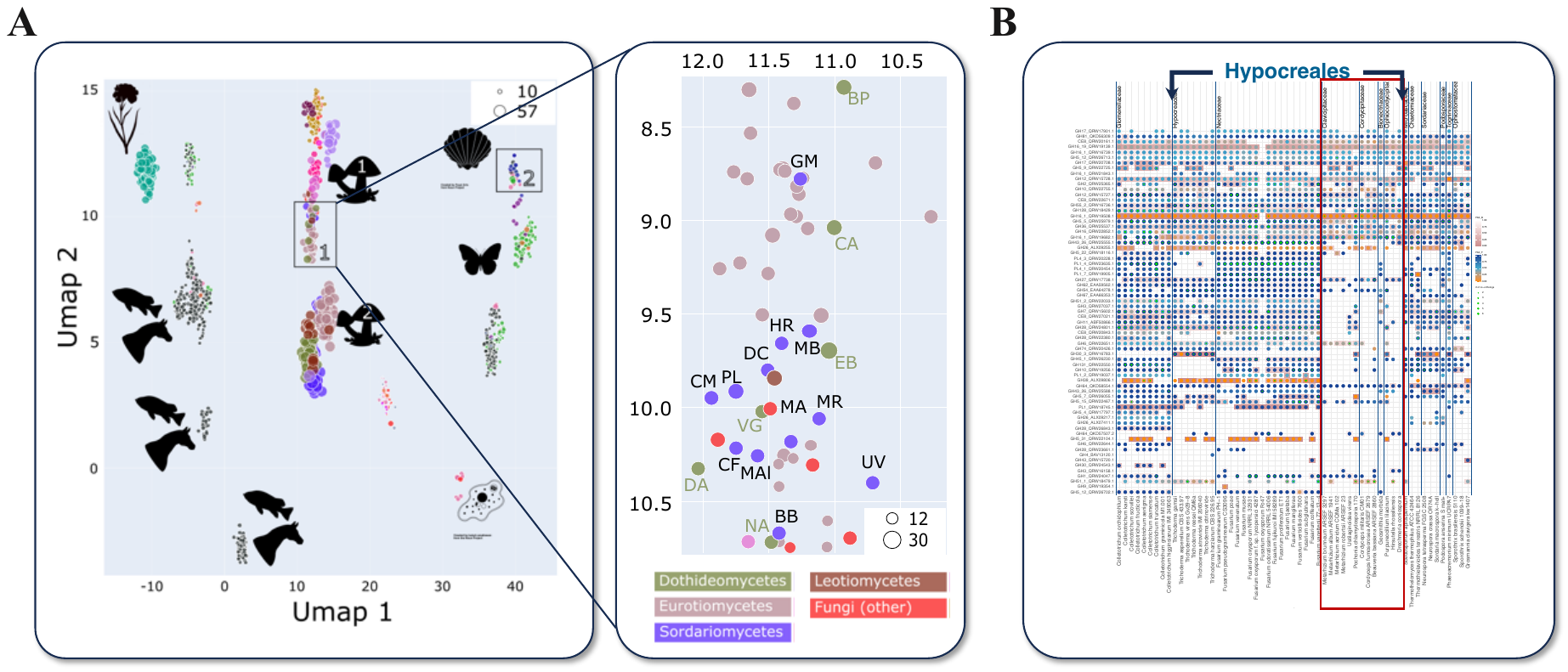

*Figure S15. fDOG reveals reduced pPCD repertoires in invertebrate-parasitising Hypocreales (Soradriomycetes).* ***(A)*** *UMAP summarizing the information about pPCD contents across the eukaryotic section of out taxon set. Fungal cluster 1 comprises 12 members of the Sordariomycetes with substantially smaller pPCD repertoires compared to the remaining xyz Sordariomycetes that are represented in fungal cluster 2.* ***(B)*** *Conventional taxon-gene matrix displaying the presence-absence pattern of orthologs to the pPCDs across the Sordariomycetes. Detected orthologs to the genes specified by the row label are indicated by a dot. The dot color represents the FAS score between the seed protein and its ortholog in a respective species. The color gradient from orange to blue indicates increasing FAS score. All taxa represented in fungal cluster 1 (see S14 A) are nested within the Hypocreales and are marked by a red box. Sordariomycetes (purple): GM – Geosmithia morbida; MB – Metarhizium brunneum; HR – Hirsutella rhossiliensis, DC – Drechmeria coniospora; CM – Cordyceps militaris; MR – Metarhizium robertsii; MA – Metarhizium acridum; MAl – Metarhizium album; CF – Cordyceps fumosorosea; UV – Ustilaginoidea virens; BB – Beauveria bassiana; PL – Purpureocillium lilacinum.*

*Figure S16. Phylogenetic profile of R. solani GH1 family hydrolase QRW24047.* ***(A)*** *Presence/absence pattern of QRW24047 orthologs across the eukaryota. Taxa have been summarized on the class level. Each dot indicates that an ortholog was found in at least 1 species subsumed in the corresponding class. The dot size indicates the fraction of species in this class with at least one ortholog. Dot color and cell color indicates the maximal FAS_F and FAS_R scores, respectively, across all subsumed orthologs.* ***(B)*** *QRW24047 is represented by several co-orthologous sequences in both human (3) and Arabidopsis (10) protein sets.* ***(C)*** *Annotations of human and arabidopsis co-orthologs with the highest feature architecture similarity to the R. solani protein.*

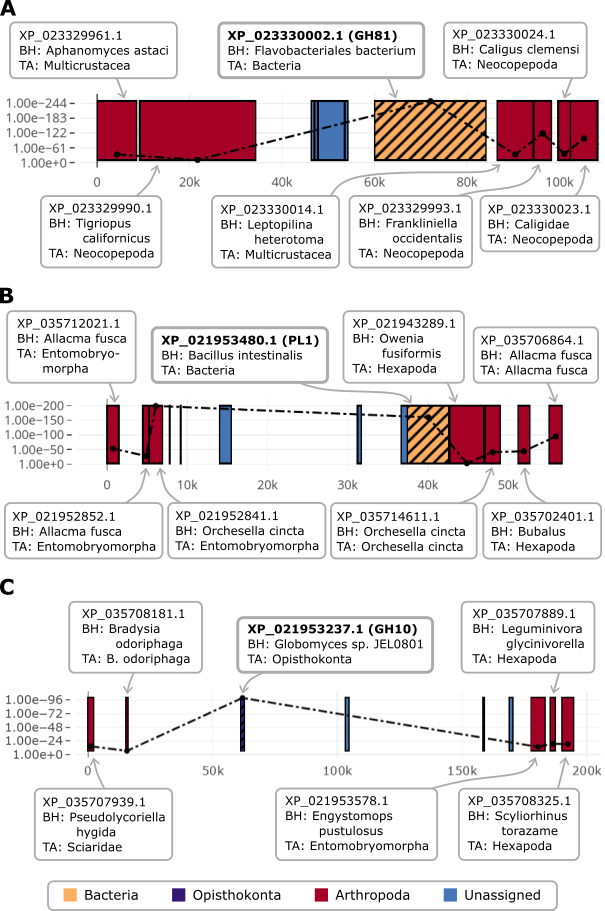

*Figure S17. Gene neighborhood analysis of plant cell wall degrading enzymes with deviant taxonomic assignments.* ***(A)*** *The Eurytemora affinis orthlogs to the fungal GH81 family endo-1,3(4)-beta-glucanase.* ***(B)*** *The Folsomia candida ortholog to the fungal PL1.* ***(C)*** *The F. candida ortholog to fungal GH10 family glyco-hydrolase. All three genes are assigned to non-animal taxa. However, they are embedded into a genomic context comprising genes whose taxonomic assignment agrees with the source organism suggesting that the carbohydrate-active enzymes have been horizontally acquired. The Y-axis represents the E-value of the best Diamond hit. BH – Best hit; TA – taxonomic assignemt. All protein identifiers represent NCBI RefSeq Accessions.*

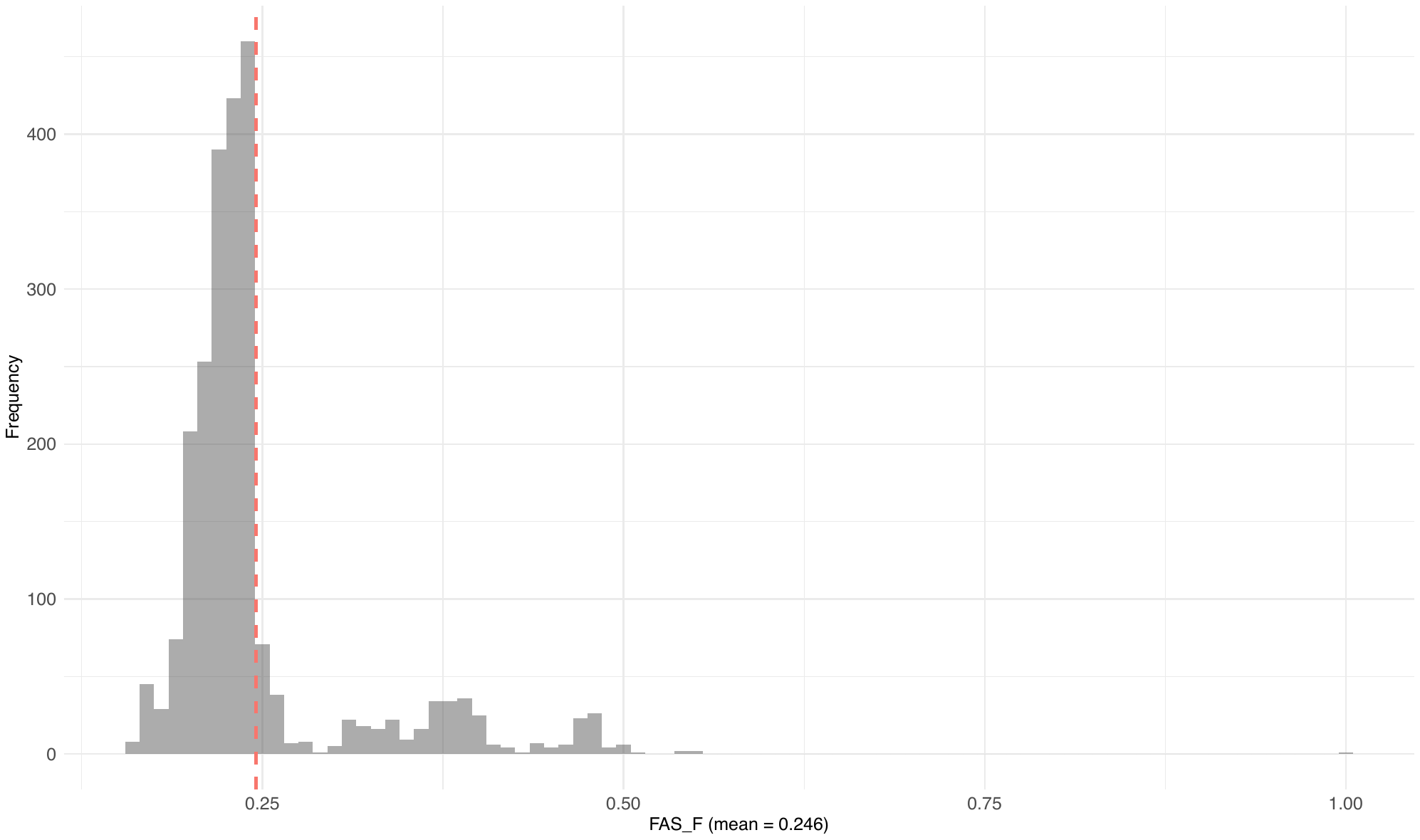

*Figure S18. Distribution of the FAS forward score for orthologs of the GH51 family protein QRW18479 of R. solani.* *The representative with a FAS forward score of 1 is the comparison of the R. solani protein against itself. See Figure 5A from the main text for the feature architecture of QRW18479.*

#### Supplementary Tables

*Table S1. Topology test result for 20 core ortholog groups that comprise only fDOG-only orthologs.* *See file SupplementaryTables-S1-S8.xlsx*

*Table S2. pPCD collection considered in this analysis.* *See file SupplementaryTables-S1-S8.xlsx*

*Table S3. Repertoires of pPCD orthologs in animals. See file SupplementaryTables-S1-S8.xlsx*

*Table S4. Taxonomic assignment of pPCD orthologs detected in Spodoptera litura. See file SupplementaryTables-S1-S8.xlsx*

*Table S5. Taxonomic assignment of pPCD orthologs detected in Eurytemora affinis. See file SupplementaryTables-S1-S8.xlsx*

*Table S6. Taxonomic assignment of pPCD orthologs detected in Bradysia coprophila. See file SupplementaryTables-S1-S8.xlsx*

*Table S7. Taxonomic assignment of pPCD orthologs detected in Folsomia candida. See file SupplementaryTables-S1-S8.xlsx*

*Table S8. Feature architecture similarities of pPCD orthologs in E. affinis, B. coprophila, and F. candida. See file SupplementaryTables-S1-S8.xlsx*
